# Intra-prostatic tumour evolution, steps in metastatic spread and histogenomic associations revealed by integration of multi-region whole genome sequencing with histopathological features

**DOI:** 10.1101/2023.02.27.530113

**Authors:** Srinivasa Rao, Clare Verrill, Lucia Cerundolo, Nasullah Khalid Alham, Zeynep Kaya, Miriam O’Hanlon, Alicia Hayes, Adam Lambert, Martha James, Iain D. C. Tullis, Jane Niederer, Shelagh Lovell, Altan Omer, Francisco Lopez, Tom Leslie, Francesca Buffa, Richard J. Bryant, Alastair D. Lamb, Boris Vojnovic, David C. Wedge, Ian G. Mills, Dan J. Woodcock, Ian Tomlinson, Freddie C. Hamdy

**Affiliations:** Nuffield Department of Surgical Sciences, University of Oxford; Nuffield Department of Orthopaedics, Rheumatology and Musculoskeletal Sciences, University of Oxford; Department of Oncology, University of Oxford; Manchester Cancer Research Centre, University of Manchester

## Abstract

Extension of prostate cancer beyond the primary site into the surrounding organs by local invasion or nodal metastasis is associated with poor prognosis. The emergence and evolution of cancer clones at this early stage of expansion and spread has not been studied in detail. We performed whole genome sequencing on 42 prostate cancer samples from the prostate, seminal vesicles and regional lymph nodes of five treatment-naive patients with locally advanced disease who underwent radical prostatectomy. Using cancer cell fractions computed from single nucleotide variants and copy number alterations, we reconstructed the tumour phylogenies, which in turn allowed us to infer key molecular steps in the progression of prostate cancer in these individuals. We mapped the clonal composition of cancer sampled across the prostate in each individual and inferred the routes of spread of cancer cells within the prostate and to seminal vesicles and lymph nodes. Based on these data, we delineated the route of tumour progression and metastasis following the transformation of adenocarcinoma to amphicrine morphology, the molecular events leading to whole genome duplication associated with a single clonal expansion and identified putative driver events associated with local invasion and lymph node metastasis. We also correlated genomic changes associated with differences in morphology and identified putative driver events associated with spread to seminal vesicle invasion and lymph node metastasis. Taken together, these findings have implications for diagnosis and risk stratification, in addition to providing a rationale for further studies to characterise the genetic changes associated with morphological transformation. Our results demonstrate the value of integrating multi-region sequencing with histopathological data to study tumour evolution and identify mechanisms of prostate cancer spread.

## BACKGROUND

Prostate cancer is a complex disease with different clinical outcomes depending on the spread of the tumour to local and distant sites. We need better risk-stratification of prostate patients to improve prognostic accuracy and enhance therapeutic strategies. Histopathological grading is an important predictor of prostate cancer prognosis. However, current pathological assessment consists of subjective morphological observations made by one or more pathologists. The limitations of the Gleason scoring system are well documented^1–4^ and patients with the same Gleason grades can have markedly differing outcomes, especially those with intermediate ISUP Grade Group 2 and 3 disease^5^. Histopathological assessment alone is incompletely predictive and prognostic, and there is an increasing need for multimodal assessment of prostate cancer.

Prostate cancer is characterised by extensive intra-tumoral heterogeneity^6^ with evidence of varying molecular expression^7^ and mutational profiles^8^ in different regions of the tumour. Recent genomic studies have detailed the evolutionary trajectories of metastatic prostate cancer and shown the utility of multi-region sampling to identify molecular features that underlie intra-tumour heterogeneity^8–11^. However, the genomic changes that occur as cancer spreads through the prostate and during local invasion and early metastasis, and their relation to histopathology remain poorly understood.

In this study, we analysed five treatment naive patients with locally advanced prostate cancer and no evidence of distant metastases. We systematically sampled tumour tissue from the prostate, seminal vesicles and local lymph nodes and used whole genome sequencing to reconstruct the tumour phylogeny in these individuals. By integrating detailed histopathological and spatial information with genomic data, we identified genomic alterations that are associated with location and changes in tissue morphology.

## RESULTS

### Multi-region sampling strategy and data integration

We performed whole genome sequencing on fresh-frozen tissue samples of the prostate, seminal vesicles and lymph nodes (Table 1, Fig 1A) to a median tumour depth of 88X using the Illumina platform (see methods for details). We systematically collected multiple samples as punches from the prostate in all cases based on histopathological evidence of tumour and from the seminal vesicles and lymph nodes where possible. We estimated cancer cell purities at 10-90% using the Battenberg algorithm (see methods). We also systematically sampled histologically normal prostate tissue punches in some patients to check for the presence of pre-malignant genomic changes. In some cases, in order to present a more complete picture of tumour spread, we augmented the data from fresh frozen samples by whole genome sequencing of histopathologically-guided sampling of formalin fixed paraffin-embedded tissue from the diagnostic archive.

**Table 1:** Summary of clinical characteristics and genomic findings.

| Patient ID | Serum PSA (ng/L) | Gleason Grade Group | pTN stage | Fresh frozen samples sequenced# | Length of follow up | Current clinical status* | Main genomic Findings |
| --- | --- | --- | --- | --- | --- | --- | --- |
| #02 | 36.8 | 5 | 3bN1 | P 8<br>SV 1<br>+ FFPE samples:<br>LN 3 | 3 years | PPP. Nodal and bone metastases. On ADT + chemotherapy (Docetaxel). | Branching evolution in the prostate. Evolution driven primarily by SNVs; polyclonal SV sample |
| #08 | 6.50 | 5 | 3bN1 | P 8<br>SV 2 | 3 years | PPP. Residual pelvic lymph node disease. On ADT + chemotherapy (Docetaxel). | Branching evolution in the prostate. Evolution driven by a number of CNAs in addition to SNVs; FOXP1 deletion in clone spreading to SV; polyclonal SV |
| #10 | 5.08 | 3 (with amphicrine differentiation) | 3bN1 | P 4<br>LN 4<br>SV 0<br>+ FFPE samples:<br>P 3<br>SV 1 | 3 years | PPP. Widespread lymph node metastases on CT. On ADT + chemotherapy (Cisplatin/Etoposide). | Branching evolution in the prostate. Amphicrine prostate cancer evolved from Adenocarcinoma; |
| #13 | 46.30 | 3 | 3bN1 | P 5<br>LN 1 | 3 years | Recurrent pelvic lymph node and bone metastases. On ADT + chemotherapy (Docetaxel). | Branching evolution in the prostate. No notable histological heterogeneity. |
| #15 | 6.83 | 5 | 3aN1 | P 2<br>LN 2 | 3 years | Salvage pelvic radiotherapy for rising PSA to 0.29. PSMA-PET negative. | Branching evolution in the prostate. Whole genome doubling;<br>chromoplexy;<br>increased SBS3; |

(UICC 8th ed)
(P=prostate, LN=lymph node, SV=seminal vesicle, ADT=Androgen Deprivation Therapy, PPP=Persistent post-operative Prostate Specific Antigen (PSA), CT=Computerised Tomography, PSMA=Prostate Specific Membrane Antigen)
\*As of February 2023

**Fig 1.**
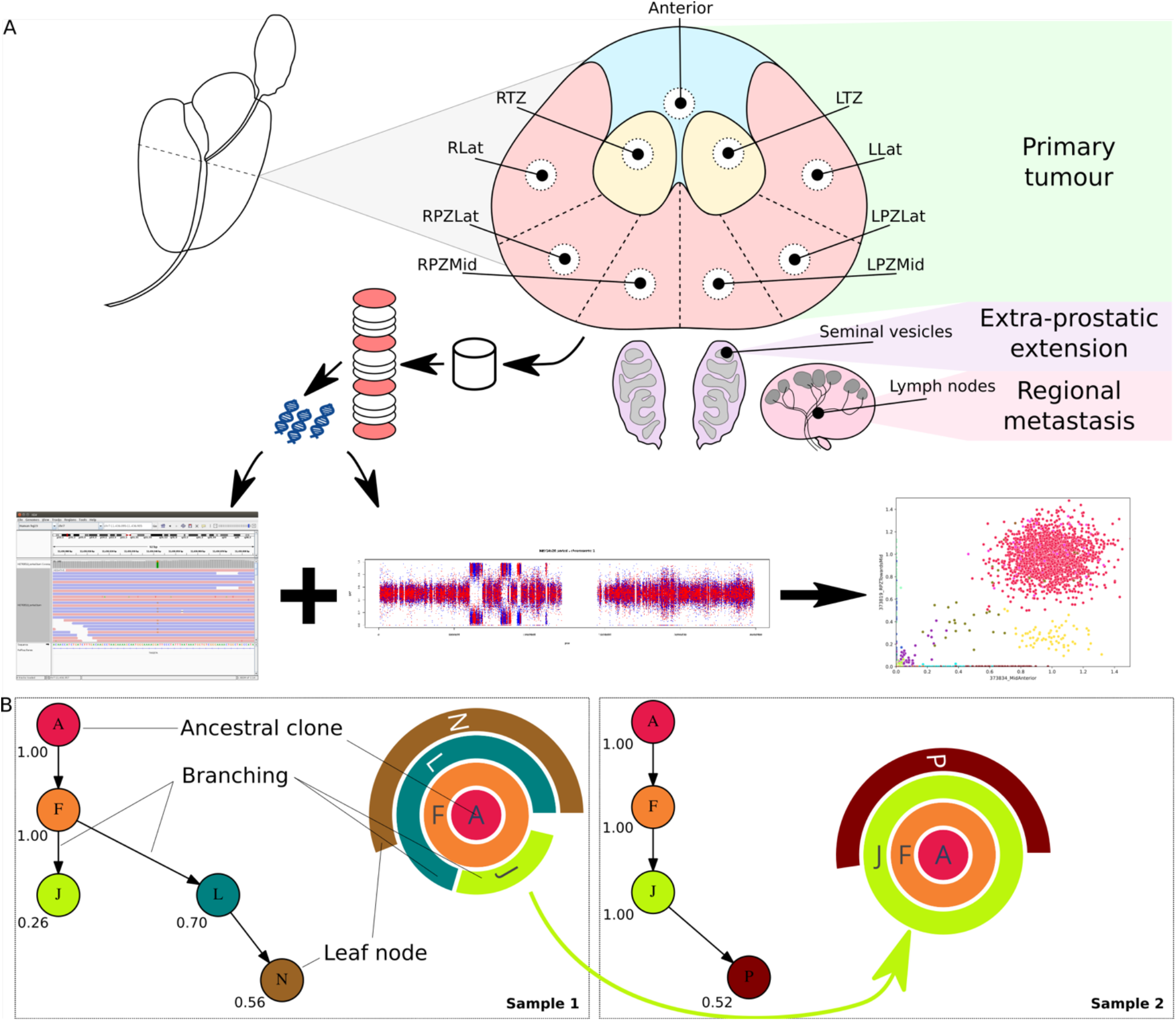
A) Schematic of sample biobanking and processing: A transverse section of the prostate was systematically sampled as 5mm diameter x 5mm thick punches to obtain samples from different regions of the prostate. In addition, samples were collected from the seminal vesicles and local lymph nodes. Punches histologically confirmed to contain tumour cells by H&E staining of intermittent sections, selected histologically normal regions, seminal vesicle and lymph node samples were sequenced over the whole genome. SNVs and CNAs calculated from the WGS data were used to infer the phylogenetic structure of the tumours. B) Depiction of phylogenetic trees as sunburst plots and determination of evolutionary routes: subclones identified in a sample are depicted as concentric circles/arcs, with the ancestral clone at the centre and each subsequent outer level representing a daughter clone. Multiple subclones at a given level indicate branches on the phylogenetic tree and hence that they are subclonal (cancer cell fraction < 1). Given two adjacent samples, e.g. sample 1 and sample 2, the tumour in sample 1 is inferred to give rise to sample 2 as the cancer cell fraction of clone J increased from 0.26 in the former to 1.00 in the latter.

Using these spatially distinct samples, we built phylogenetic trees based on somatic small nucleotide variants (SNVs) and copy number alterations (CNAs) to assess cancer evolution in each patient. In addition to the overall phylogenetic tree for each patient, we inferred inter-sample clonal relationships based on the median cancer cell fraction of the SNVs in each clone and the spatial context of each sample. We depict the clonal composition of each sample as a sunburst plot (Fig 1B) and the inter-sample relationships were superimposed on the histology images of the whole prostate to construct a ‘clone map’ for each patient. We present these results below.

A summary of the somatic mutations in the sequenced samples revealed that *PTEN* is deleted in nearly all samples across the 5 patients (Fig 2). Further, tumour suppressor (*TP53, RB1*) and DNA repair (*BRCA2*) genes were altered in more than one individual. SNVs and CNAs were commonly present across several samples in each individual, pointing to a shared evolutionary history. Samples from #15 had the highest number of mutations affecting coding regions. We also found a higher proportion of C>T mutations in one of the four samples from #15.

**Fig 2.**
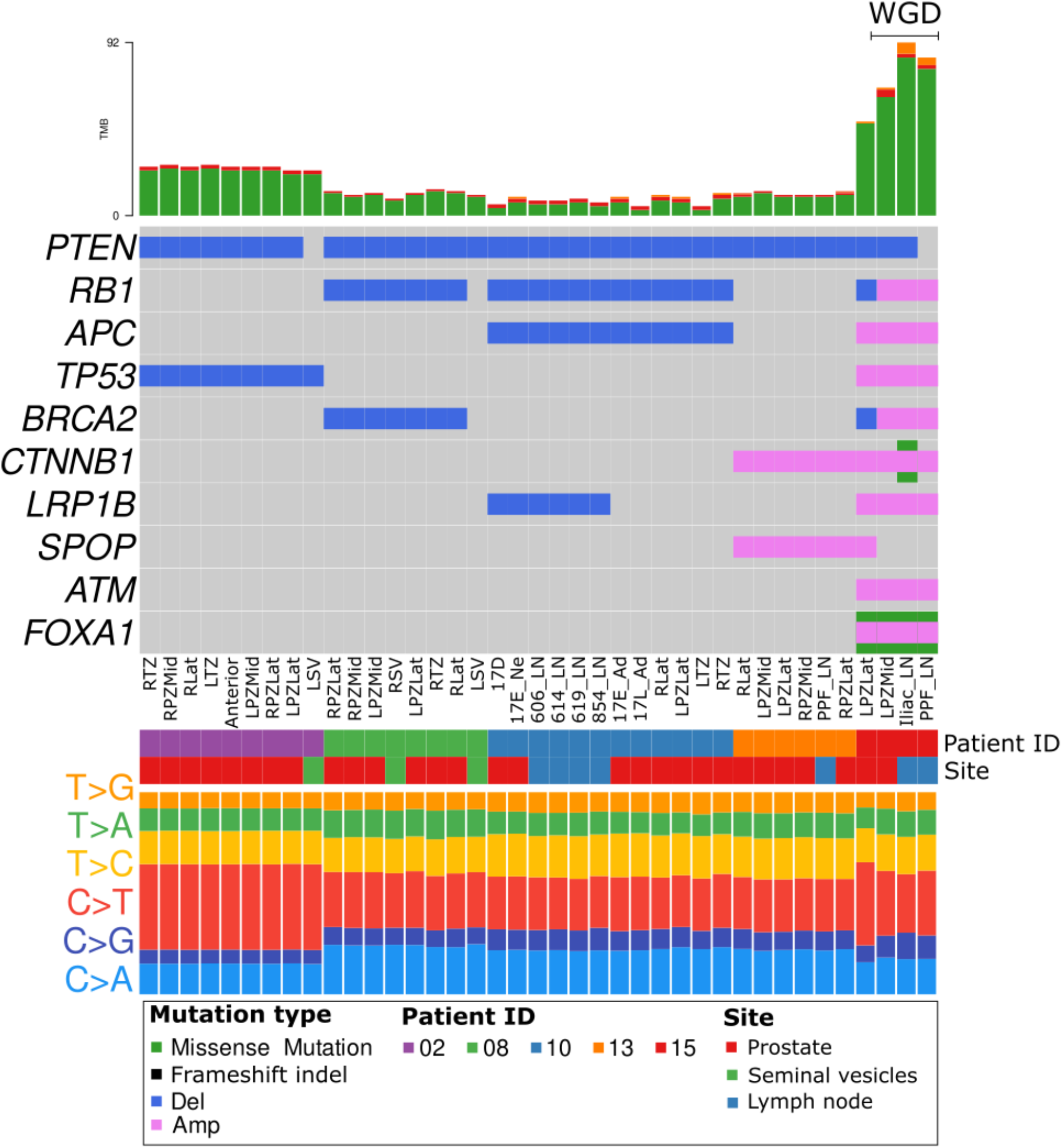
Oncoplot summary of SNVs and CNAs affecting known prostate cancer diver genes (Intogen) in the sequenced samples. Samples are grouped by patient ID and annotated for sample site. Integer copy numbers < 2 and > 2 were classified as deletion (Del) and amplification (Amp) respectively. Proportions of the six possible types of mutational conversions in the SNVs are shown at the bottom. (TMB: Total Mutation Burden; in this figure, refers to SNVs in coding regions)

### Clone maps reveal evolution and spread of cancer cells within the prostate

In patient #02 (Fig 3A), the LPZLat sample represents the earliest part of the malignancy, with origin of new clones as the cancer spreads along the peripheral zone to the right side. We found that the ancestral clone (clone A) comprising several copy number changes (e.g. 10q23 LOH, 16q22 LOH, 17p13 LOH) eventually gave rise to a number of subclones that were exclusively distinguished by SNVs with no further changes observed in copy number. The cancer in the seminal vesicle sample (LSV) was determined to be polyclonal (defined as a sample consisting of two or more subclones sharing a parent clone), suggesting that this is a result of either multi-clonal invasive spread or seeding from multiple metastatic events. Lymph node metastases arose from Clone A (5A_LN) and Clone J (6A_LN, 7A_LN). Clone J was also the source of the majority of the clonal heterogeneity in this patient, as it gave rise to multiple independent daughter clones (E, I, O, P). However, no copy number changes or SNVs affecting coding genes could be attributed to this clone.

**Fig 3.**
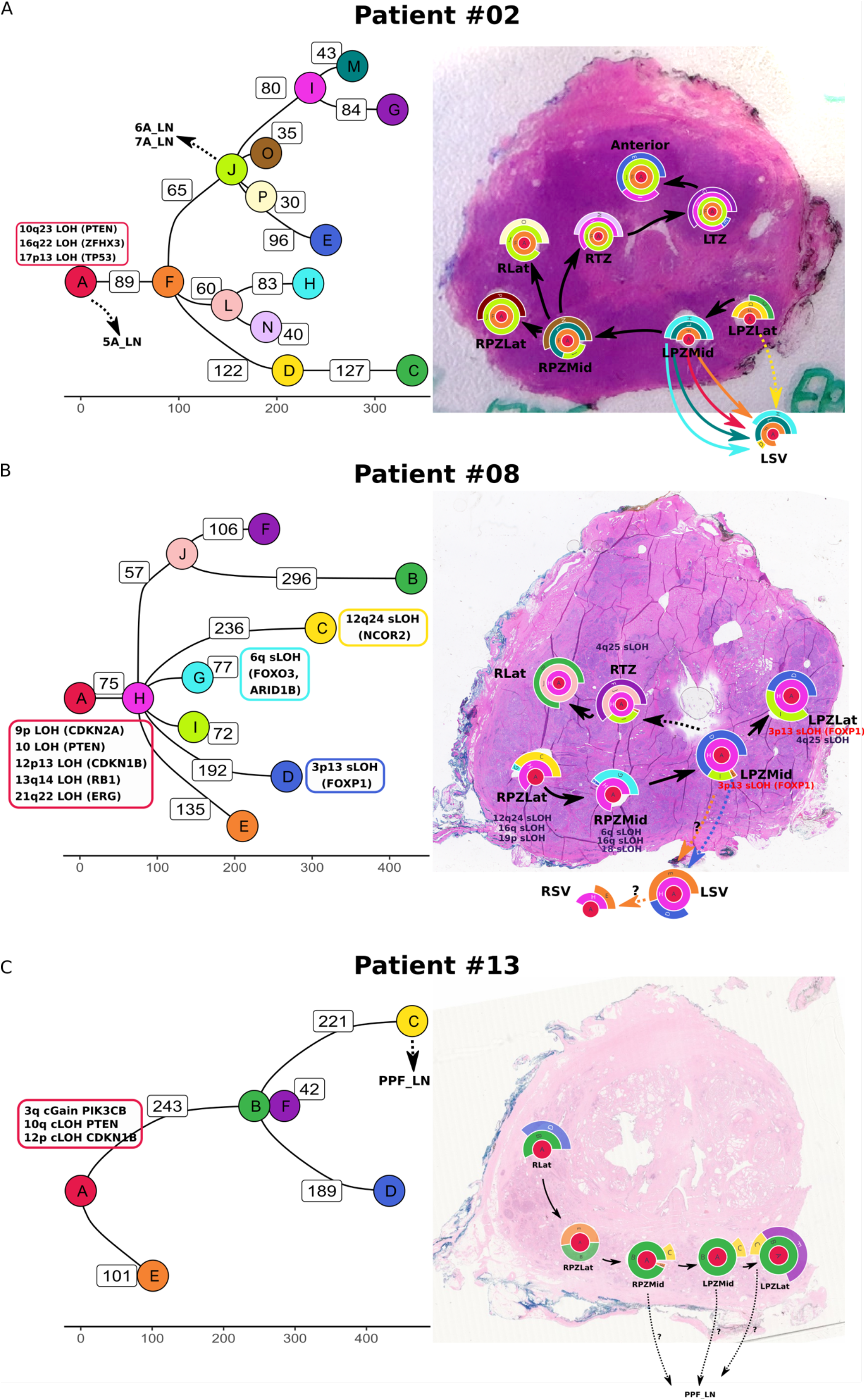
A) Phylogenetic trees and clone maps of patient #02, B) patient #08, C) patient #13, and D) patient #03, showing diverse routes of tumour evolution within the prostate, local invasion to seminal vesicles, and metastasis to lymph nodes. The phylogenetic trees are annotated with SNVs and CNAs involving known prostate cancer driver genes (Intogen). The numbers on the edges of the phylogenetic tree and the X-axis scale represent the number of SNVs assigned to the daughter clone. Metastatic events to lymph nodes are annotated on the phylogenetic tree with dashed arrows. Clone names are sorted based on cluster size, i.e., A > B > C and so on.

In patient #08 (Fig 3B), the earliest tumour location was inferred to be RPZLat and it spread along the peripheral zone to the left. The ancestral clone harboured copy number alterations involving several known driver genes (9p LOH, 10 LOH, 12p13 LOH, 13q14 LOH, 21q22 LOH). Several subclonal copy number alterations were also prominent in the leaf nodes (tips) of the evolutionary tree, e.g. 12q24 LOH in clone C, and 6q LOH in clone G, affecting known prostate cancer driver genes (Intogen) such as *NCOR2, FOXO3* and *ARID1B*. Clone D in particular harboured a loss of *FOXP1* (3p13 LOH), a known tumour suppressor gene in prostate cancer^12^, and notably this clone was present in a seminal vesicle sample (LSV). Knockdown of *FOXP1* in LNCaP prostate cancer cells resulted in increased migration in vitro (Supplementary fig. 7). Clone H is the source of the majority of clonal heterogeneity in this patient, and is the source of 6 subclones. However, no protein-coding mutations could be attributed to this clone. As in patient #02, the tumour in the seminal vesicles is polyclonal, again suggesting an invasive spread from the prostate.

In patient #13 (Fig 3C), RLat represents the earliest region of the tumour, with the tumour spreading along the peripheral zone towards the left. Clone B is the last dominant clone (present in all intra-prostatic samples) and was the source of 3 subclones (C, D, F). Lymph node metastasis (PPF_LN) was determined to arise from clone C, although low tumour purity in the lymph node sample precluded a more detailed analysis.

### Amphicrine prostate cancer arises from adenocarcinoma

In patient #10, histology of the lymph nodes revealed cancer in 12/22 nodes on the right side and 1/13 nodes on the left side. All nodal disease was amphicrine morphology (Fig 4A). Of these, four lymph nodes were biobanked and sequenced - amphicrine morphology which was not seen in the intra-prostatic fresh-frozen samples (which were adenocarcinomas histologically). Amphicrine cancer is a recently described variant of prostate cancer on the spectrum of neuroendocrine differentiation, where primary cancers display features of both exocrine (classic acinar adenocarcinoma) and neuroendocrine prostate cancer^13,14^. On standard diagnostic staining, the primary prostate cancer in this case was negative for neuroendocrine markers, but the nodal disease was positive for *AR* and markers of NE differentiation (Synatophysin, and CD56) (Figure 4A) with a Ki67 index of 60%. Most of the intra-prostatic malignant tumour was present towards the base of the prostate close to the seminal vesicles. Diagnostic H&E sections from this area revealed the presence of adenocarcinoma and amphicrine carcinoma in the prostate close to and in the base of the seminal vesicles with coexistence of the two histological subtypes (Fig 4C).

**Fig 4.**
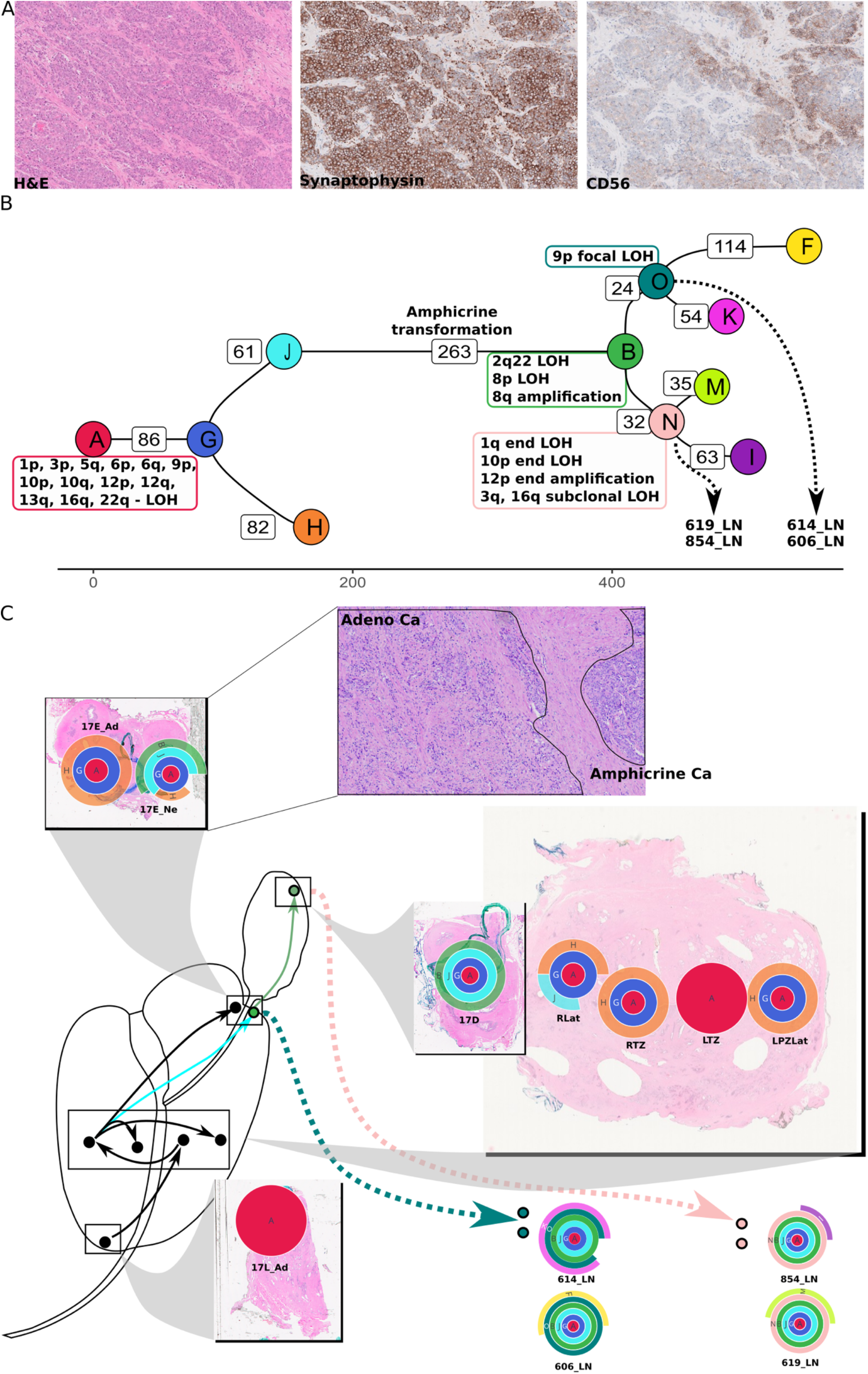
A) Amphicrine morphology in lymph nodes, as seen by H&E staining, immunohistochemistry for Synaptophysin and CD56. B) Phylogenetic tree for patient #10, with several copy number changes at the branching points. Clones that were likely sources of metastatic seeding are connected to the lymph node sample IDs by dashed arrows. D) Clone map showing inferred route of cancer progression; clone J in the RLat punch expands in 17E_Ne and is associated with transformation from adenocarcinoma to amphicrine appearance.

After histology-guided dissection of FFPE sections from the amphicrine and adenocarcinoma regions separately, extracted DNA was sequenced to enable a detailed reconstruction of tumour evolution in this patient. This analysis revealed that the adenocarcinoma transdifferentiated into amphicrine morphology during its spread through the prostate, with prominent genomic changes including the loss of 8p, amplification of 8q, and loss of 2q22 (Supplemental information section 5). Two seeding events were inferred, leading to metastatic growth in 614_LN/606_LN and 619_LN/854_LN lymph nodes respectively, based on SNV clustering as well as copy number analysis (Fig 4B and 4C). The first seeding event to 614_LN/606_LN was inferred from the presence of a focal LOH in 9p23 (affecting the PTPRD gene); this focal LOH was not detected in the other lymph node or intra-prostatic samples. Hence, this CNA was determined to occur prior to or coincident with the emergence of clone O. Due to the features shared with 17E_Ne, and the absence of features from 17D, the seeding to 614_LN/606_LN was inferred to be between these two prostate/seminal vesicle samples. The second seeding event to 619_LN/854_LN lymph nodes was inferred by the presence of a shared CNA with 17D (12p end amplification), placing its origin chronologically after 17D. Several copy number changes (1q end LOH, 10p end LOH, 3q and 16q subclonal LOH) are also shared by 619_LN/854_LN lymph nodes, suggesting that they were coincident with the emergence of clone N. The clone map for this patient uncovered a complex 3-dimensional evolutionary history, with lymph node metastases arising from or near the seminal vesicles. Copy number analysis revealed telomeric allele imbalance in chromosomes 1q (LOH), 6p (LOH), 8p (LOH) and 12p (gain). In addition, a sample (Anterior punch) annotated as benign by histopathological assessment, but close to the main tumour was found to harbour a mutant clone with ∼250 SNVs of a lineage distinct from the main tumour. No prostate cancer-specific driver mutations could be identified to explain the emergence of this clone and no copy number changes were detected.

### Distinct histopathological features and multiple metastatic seeding events in patient #15

In patient #15, from whom two intra-prostatic and two lymph node samples were analysed, we were able to reconstruct the phylogenetic tree from a complex set of chromosomal gains and losses. *PTEN* loss and *TP53* frameshift insertion were observed in all samples from this patient. From the clonal composition and the phylogenetic tree (Fig 5A, 5B), we inferred that clone H originated in LPZLat and expanded to become clonal in LPZMid sample. This clone eventually seeded metastasis to both the lymph nodes (Iliac_LN and PPF_LN) through its daughter clone A. In addition, two additional seeding events to the PPF_LN could be inferred, one event from LPZMid (clone I) and another from LPZLat (clone G). The relative timing of these seeding events could not be determined from the data.

**Fig 5.**
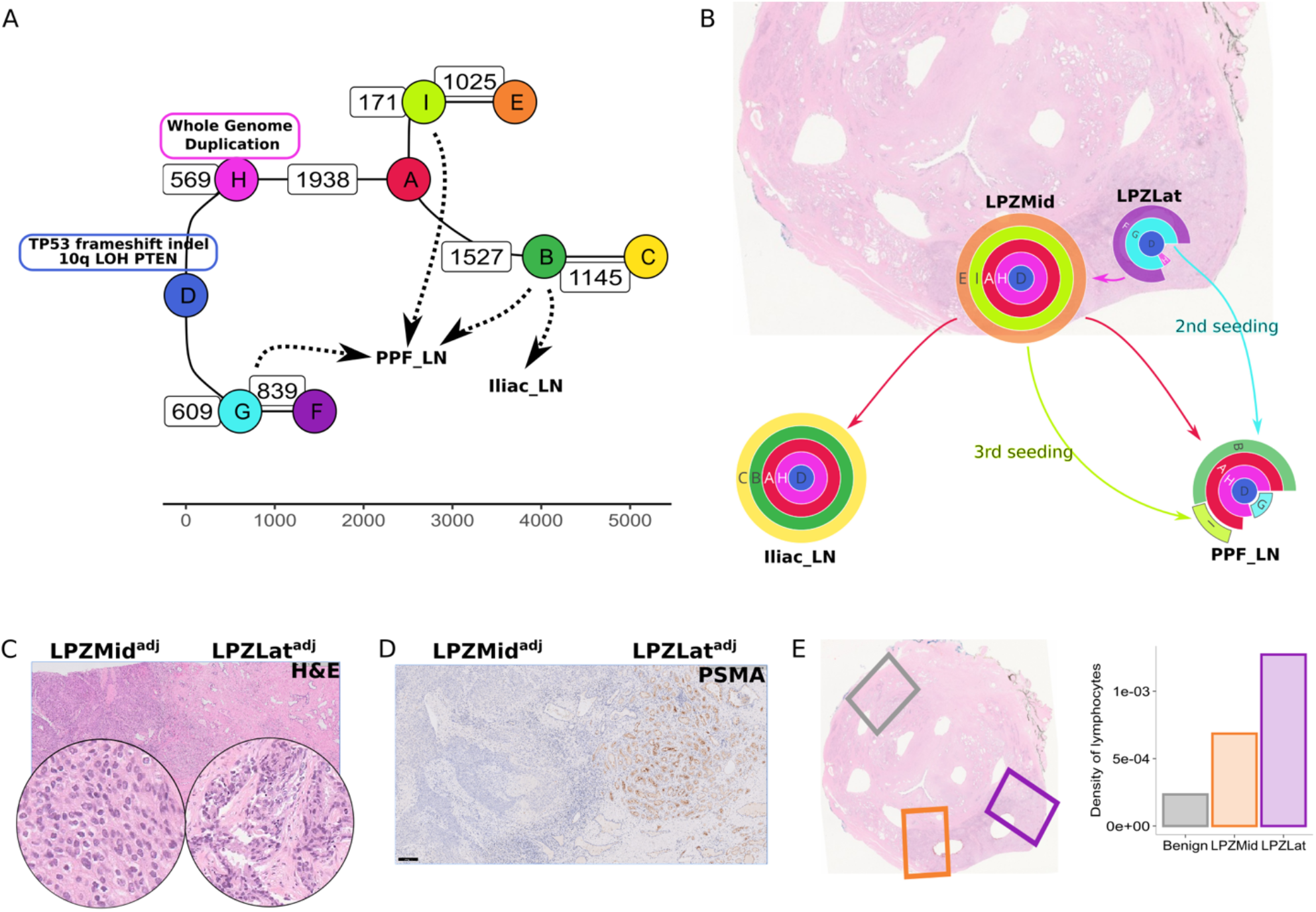
A) Phylogenetic tree for #15 showing clones I, G and C as potential sources of metastatic seeding to lymph nodes B) Clone map of patient #15 showing routes of intra-prostatic spread and metastasis to lymph nodes. Clone H (pink) expands to become clonal in LPZMid. C) A H&E section adjacent to the sequenced samples with areas corresponding to LPZMid and LPZLat on a plane close to the punches (superscripted with *adj* to denote this) showing different morphologies with D) matching PSMA expression and E) numbers of tumour infiltrating lymphocyte counted by image analysis in corresponding regions compared to a distant non-tumour region.

In histopathological analysis, the two intra-prostatic regions corresponding to the sequenced samples had a distinctly different morphology, with a clear demarcation between the two (Fig 5C). The LPZMid sample had a higher Gleason score of 10 (5+5)/Grade Group 5, versus LPZLat which had a Gleason Score of 7 (4+3)/Grade Group 3. There was also extensive solid pattern intraductal carcinoma adjacent to the LPZMid punch, but not LPZLat. This difference between these two samples was also reflected in the PSMA staining pattern, with minimal or absent staining in the Gleason Score 10 areas around LPZTowardsMid (Fig 5D) (H Score 10) but retained in the Gleason Score 7 areas around LPZLat (H Score 100). The cancer metastases in the lymph nodes matched the morphology in LPZMid. There was also a high distribution of tumour infiltrating lymphocytes in the LPZLat sample, whereas TILs were notably less abundant in LPZMid (Fig 5E).

### Whole genome duplication and extensive chromoplexy distinguish the intra-prostatic samples in #15

In association with the histological transformation observed in patient #15, we identified whole genome doubling (defined as ploidy > 3) in three out of the four samples (Fig 6A). In the sample without whole genome doubling (LPZLat), the 84% of the genome was subclonal (as opposed to 18% subclonality in LPZMid), suggesting that this sample is composed of a mixture of clones with differing copy number profiles. *PTEN* was hemizygous in this sample as seen by positive IHC staining (Fig 6B) and copy number profiling (Fig 5C), in contrast to LPZMid where PTEN was completely deleted. Analysis of structural variants revealed several inter-chromosomal translocation events (Fig 6C) indicating chromoplexy and extensive chromosomal fragmentation suggestive of genomic instability in LPZLat. While several structural variants are shared between LPZLat and LPZMid (corroborating the evolutionary origin of the latter from the former), the number of inter-chromosomal translocations is higher in the former (60 vs 48). Translocations present in the PPF_LN and LPZLat samples (chr5-chr19, chr13-chr15, chr2-chr5) but not in the LPZMid or Iliac_LN samples, corroborates the additional metastatic seeding event (clone G) from LPZLat to PPF_LN as inferred from SNV clustering (Fig 5A).

**Fig 6.**
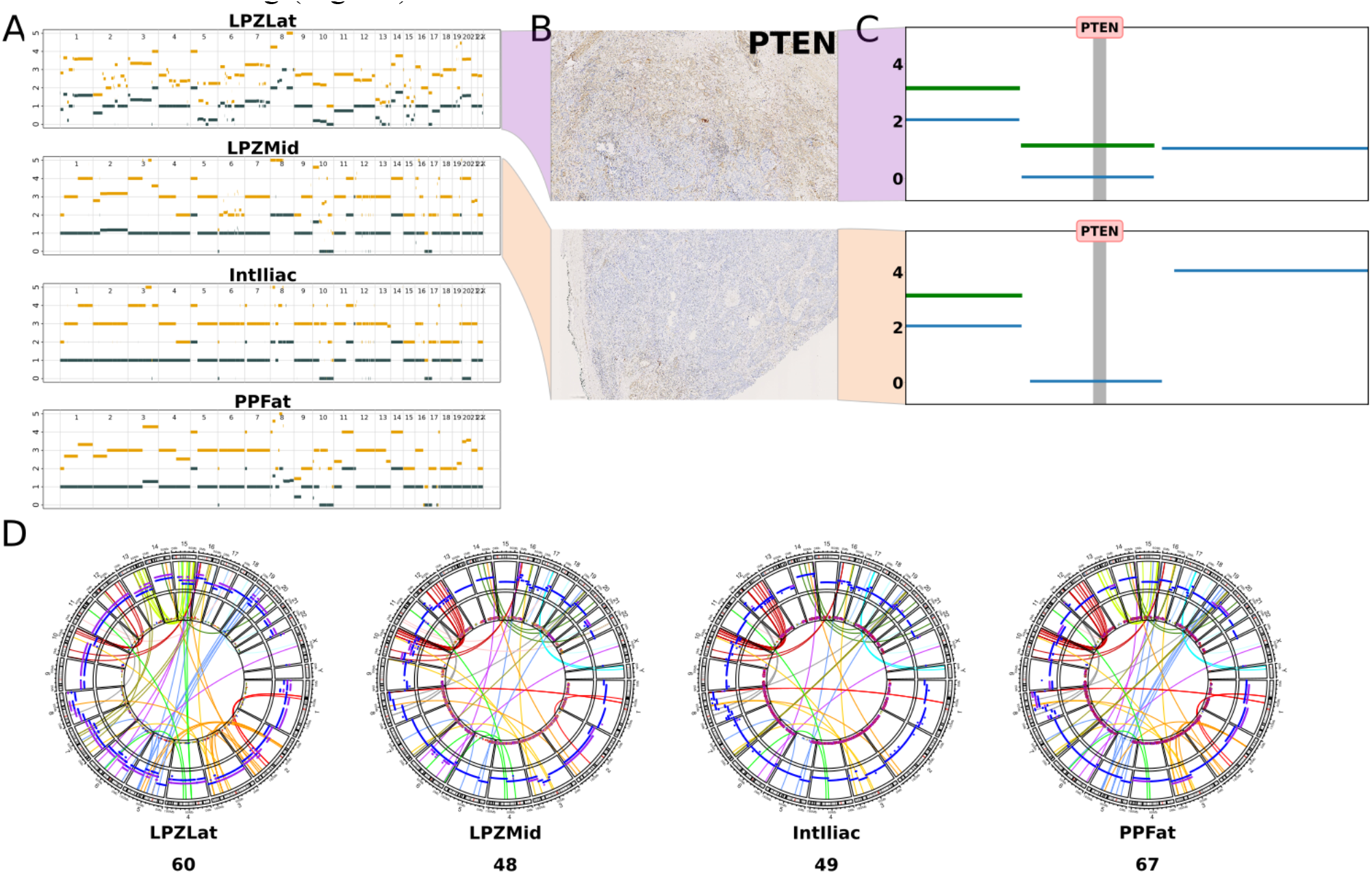
A) Copy number plots for #15 generated by Battenberg showing a subclonal profile for LPZLat and whole genome duplication in LPZMid, IntIliac and PPFat samples. B) PTEN immunohistochemistry staining corresponding to LPZLat (top) and LPZLatMid (bottom) samples and C) their corresponding PTEN copy number profiles. D) Structural variants plotted as circos plots, with different coloured links based on the chromosomes in which the variants’ start coordinates are located.

### Phylogenetic tree branches in #15 show differences in mutational signatures

Unlike other individuals, the truncal or ancestral clone in #15 was not the clone with the highest number of mutations (Supplemental information section 6). Hence, we hypothesised that a new mutational process could have led to an increased mutation rate later in the phylogenetic tree. To investigate this further, we subdivided the SNVs into five groups based on the four sub-sections of the phylogenetic tree (GF, HA, BC, IE) and one truncal clone (D) and analysed their trinucleotide mutational signatures (Fig 7A). While C>T mutations were the most abundant type in D and GF, this was not the case in HA, BC and IE. Deconvoluting these mutational signatures into known signatures from COSMIC v3.2, we discovered that SBS3 (associated with defective homology directed repair) increasingly contributed to the overall signature in HA, BC and IE (Fig 7B). This suggests that there is a change in mutational processes starting at clone H. However, *BRCA1* and *BRCA2* genes were intact (Supplemental information section 8: CNA profiles zoomed in on *BRCA1, BRCA2, PALB2 loci*) in LPZMid and there was no evidence of increased methylation in *BRCA1* and *BRCA2* promoter regions (Supplemental information section 9: Methylation EPIC array data). This led us to explore alternative explanations for the increased contribution of SBS3 signature in HA, BC and IE subsections. We observed that the increased contribution of signature SBS3 coincided with whole genome duplication. We also observed increased *PHH3* staining in LPZMid compared to LPZLat (Fig 7C) indicating a higher number of cells in G2/M in the former region.

**Fig 7.**
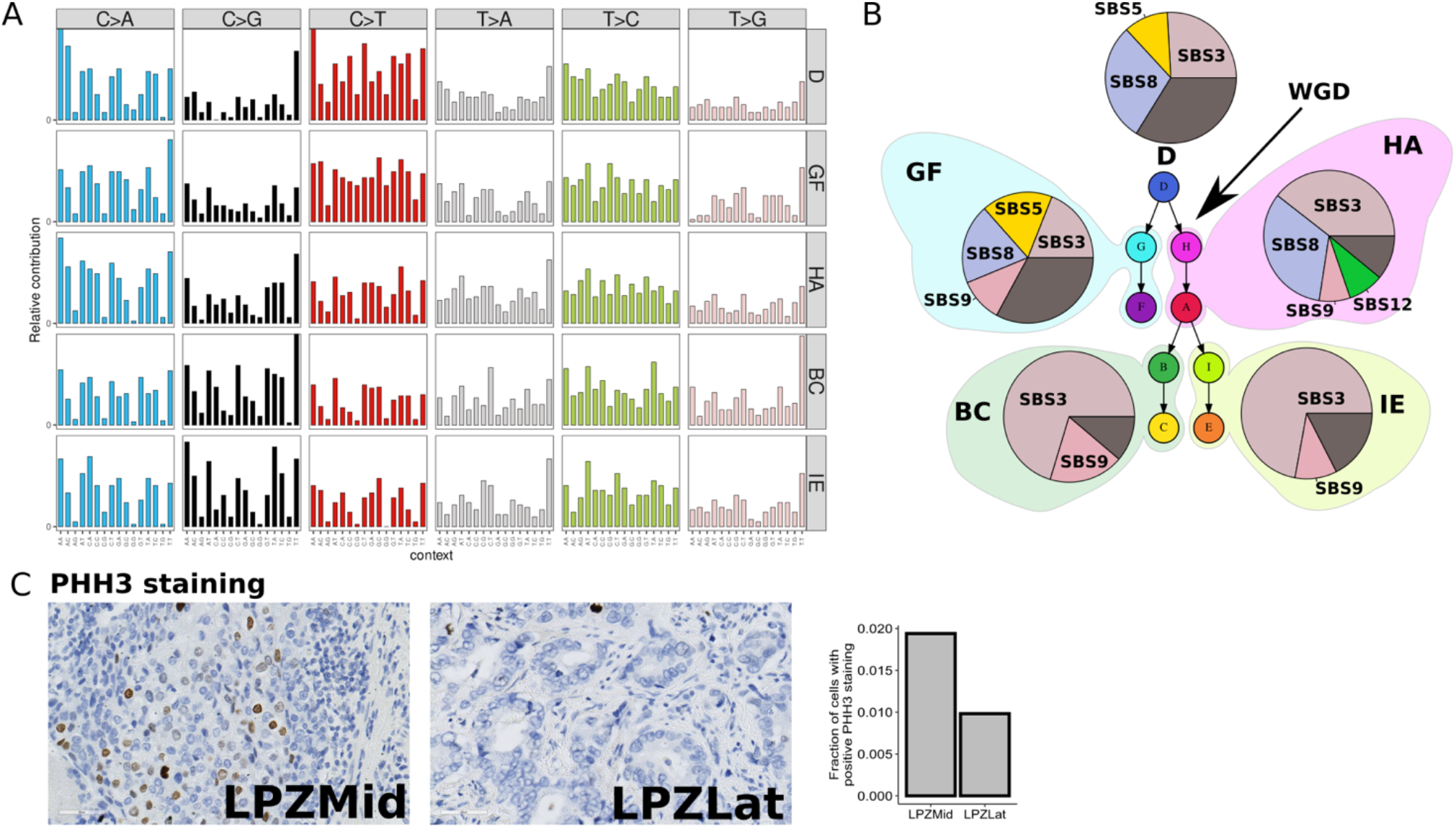
A) Trinucleotide mutational signatures in each subsection of the phylogenetic tree for patient #15 B) Pie charts showing increasing contribution of signature 3 in tree sub-sections HA, BC and IE compared to D and GF C) Immunohistochemical staining for PHH3 (positive in G2/M phase of cell cycle), plotted as the fraction of positive cells, shows a higher proportion in the tissue region corresponding to LPZMid.

### Histological differences are associated with genomic changes

In order to understand the relationship between genomic changes and histology, we analysed the histological characteristics of the punches and correlated these with their corresponding clonal composition. We observed histological differences in adjacent punches in 4 patients (Fig 8). In #02, prominent intraductal carcinoma (IDC) was present in RPZMid in contrast to no IDC in RTZ. Similarly, in #08 predominantly cribriform intraductal carcinoma, Gleason grade 4+4 was present in RPZLat whereas RLat was comprised of a higher Gleason grade tumour (4+5) without cribriform morphology. In #10, adjacent regions within the same FFPE section displayed features of adenocarcinoma and amphicrine carcinoma. In #15, LPZLat consisted of a Gleason grade 4+3 tumour and the neighbouring punch LPZMid was classified as grade 5+5 with no discernible glands. In all these cases, each histological subtype was found to predominantly consist of clones from a separate branch of the phylogenetic tree. We quantified the total number of SNVs uniquely present in either sample in the pair as a measure of the evolutionary distance between the two morphologies – 223, 666, 324 and 4582 unique SNVs were seen in patients #02, #08, #10 and #15 respectively.

**Fig 8.**
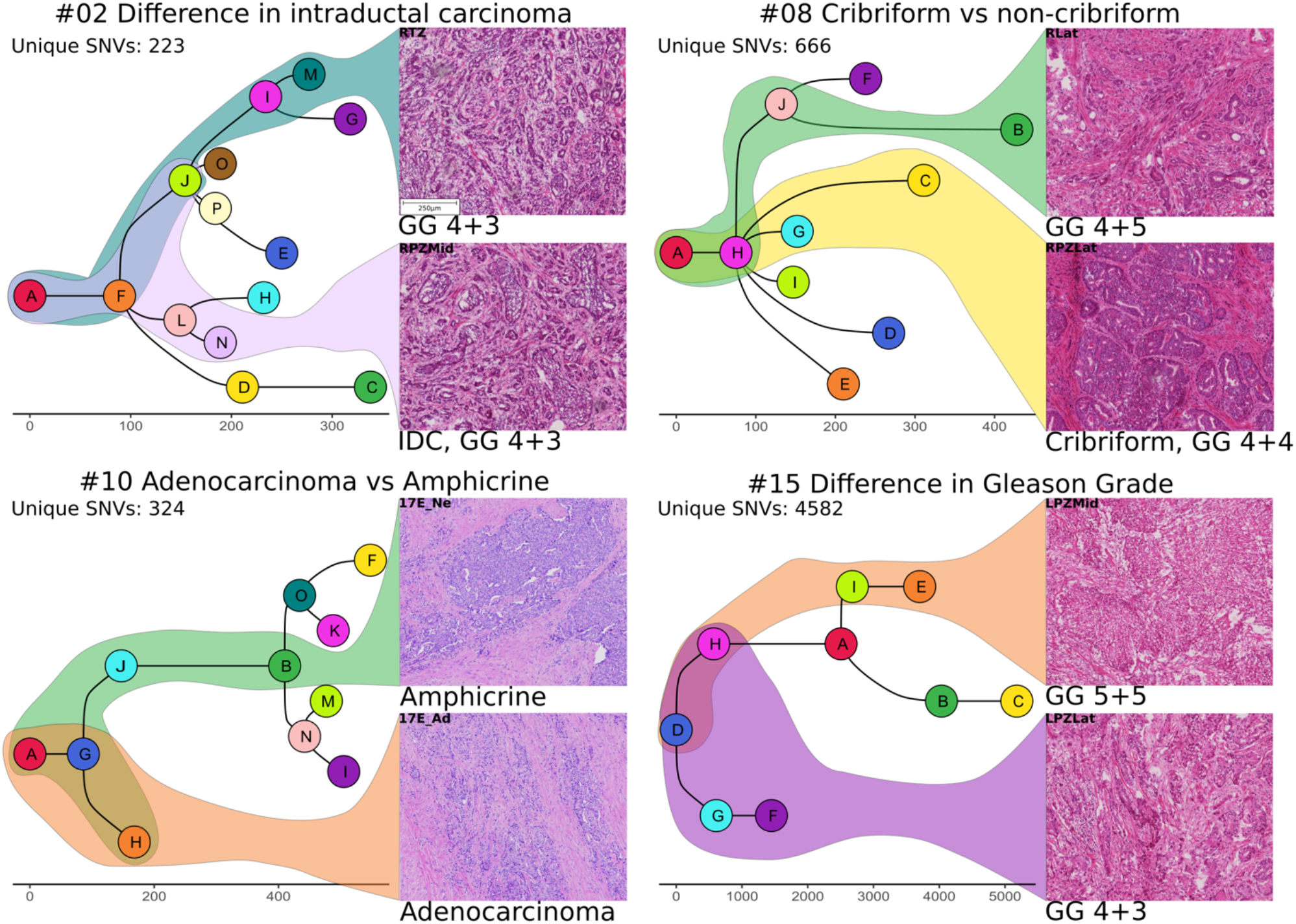
Pairs of H&E-stained sections (frozen: #02, #08, #15; FFPE: #10) corresponding to sequenced regions from each of four patients (4X magnification). Histological differences within each pair are highlighted and the corresponding branches of the phylogenetic trees are shown in the colours of the respective leaf/tip nodes. The total number of SNVs unique to either of the two histologically distinct samples are represented as “Unique SNVs”. (GG: Gleason grade; IDC: Intraductal carcinoma of the prostate)

### Other general findings

We observed a high degree of branching evolution within the prostate. In patient #02, branching occurs at nodes F, I, J and L resulting in 8 daughter clones within the prostate. In patient #08, branching at nodes H and J results in 7 daughter clones within the prostate. Intra-prostatic branching evolution was also seen to a lesser extent in #10 (at node G), #13 (at nodes A and B) and #15 (at nodes D and A).

We also observed multiple seeding events to lymph node metastases (patient #02, patient #10, patient #15). This raises the possibility that metastatic potential was acquired in a clone that is not most directly associated with a seeding event. For example, in patient #02, while metastatic seeding occurs from clone A (5A_LN) and clone J (6A_LN, 7A_LN), no SNVs were observed in the coding regions of known driver genes and no new copy number alterations were acquired between clone A and J.

In the samples analysed, extensive tumour evolution occurred in the prostate, whereas fewer subclonal clusters could be identified in lymph node samples. However, in patient #10, continued evolution occurred in the lymph nodes with each of the 4 lymph nodes harbouring its own unique subclone.

In all patients except patient #10, the earliest tumour location is in the peripheral zone and the malignancy spreads circumferentially. The patterns of inferred cancer spread within the prostate suggest a predilection for the cancer cells to be limited to the prostate capsule (although there is histological evidence of extraprostatic extension, which is classified as T3a, in limited areas in all the patients in this study). In patient #10, the earliest tumour location is in the apex of the prostate, with lymph node metastases arising from the seminal vesicles.

For the majority of intra-prostatic clones in patient #02, no specific driver mutations (copy number or SNV) could be attributed. Specific driver genes can be attributed to SV invasion in #08 and lymph node metastases in #10.

## DISCUSSION

Using a systematic approach to biobanking and sequencing analysis, we integrated genomic data with histopathological and spatial information in order to delineate the routes of tumour evolution in intra-prostatic spread, local invasion and early metastasis in treatment-naive prostate cancer patients. We reconstructed evolutionary trajectories in five individuals, generating clone maps to visualise the spread and expansion of tumour clones. This approach revealed evolution primarily driven by SNVs in patient #02, from an ancestral clone with several copy number alterations. On the other hand, several copy number changes involving known cancer drivers were observed in patient #08, with extensive branching evolution within the prostate. These cases throw new light on the source of the extensive intra-tumour heterogeneity seen within prostate cancers, building on work reported by Cooper et al.^8^, who depicted intra-tumour genomic heterogeneity using phylogenetic analysis of multi-region samples. The inclusion of seminal vesicle and lymph node samples revealed the specific clones that resulted in local invasion and metastasis.

Presence of cancer in the seminal vesicles is staged as T3b and is associated with worse prognosis than organ-confined disease. The SV samples in this study were polyclonal, suggesting invasive spread from the prostate as the most likely mechanism, corroborating the findings from Woodcock et al.^11^. However, multi-clonal seeding through metastatic spread cannot be ruled out. In patient #10, lymph node metastases arose from the base of the prostate and the seminal vesicles, which concurs with the reports that seminal vesicle invasion is associated with metastasis and worse prognosis^15^.

In patient #10, we show the genomic correlates involved in the transformation of adenocarcinoma to amphicrine morphology. Amphicrine cancer is a relatively rare subtype of prostate cancer having a distinctive morphology with synchronous dual exocrine and neuroendocrine (NE) differentiation. Morphologically, these tumours show solid/nested growth with cells with amphophilic cytoplasm, vesicular nuclei and macronuclei, without the features of small cell or large cell NE carcinoma. Immunohistochemically the tumours are positive for PSA, AR, ChromograninA or Synatophysin and have a high Ki67 index (median 50%)^13^. While data are limited due to its rarity, it is thought that amphicrine prostate cancer is often associated with metastatic disease^16^ and may be responsive to AR-targeted therapy^17^. More recent work by Graham et al. suggests that there may be differences in the clinical outcomes of treatment-naïve and post-treatment amphicrine prostate cancer^18^. We were able to map the 3-dimensional spread of cancer in patient #10 using histopathology as a guide for sample selection. The spatially proximal but morphologically and genetically distinct areas within the same tumour have very different outcomes: the adenocarcinoma did not metastasise, whereas the adjacent amphicrine cancer metastasised to several lymph nodes. The amplification of 8q and loss of heterozygosity of 8p are prominent changes associated with the amphicrine transformation in this patient. The enhanced understanding of cancer spread in 3D in this patient also shows the limitation of representing clonal spread as 2D clone maps, as this is likely to be a simplification of a more complex series of events in the other patients.

In patient #15, we identified a whole genome duplication event; subclonality (copy number, SNVs) and genomic instability were also observed in the LPZLat sample. López et al. proposed that whole genome doubling could be a mechanism for reduced genomic instability^19^. PHH3 staining is specific for the M phase of the cell cycle^20^ and hence, a higher staining in the LPZMid sample suggests a delayed cell cycle. Increased genomic complexity is thought to induce greater replication stresses^21^ and upregulation of the NHEJ repair pathway in polyploid cells^22^. The increased but error-prone repair through the NHEJ pathway may explain the SBS3 signature in conjunction with WGD in patient #15. Taking these findings together with the increased immune cell infiltration in the pre-WGD region, we propose a model of tumour progression in patient #15 whereby increased genomic instability due to inactivation of TP53 could have resulted in rapid genomic changes, with such a region acting as a ‘clone factory’ where several daughter clones were produced. The high rate of mutations also could have led to increased immune surveillance, which combined with the effect of deleterious mutations caused most daughter clones to fail to survive. WGD acted as an escape mechanism by allowing repair of DNA damage through the NHEJ pathway. Thus, even though WGD cells may have a lower rate of growth due to cell cycle delay, they were better able to survive due to better tolerance to deleterious mutations and reduced immune surveillance.

Histopathological evaluation of tissue samples is central to prostate cancer diagnosis. Several morphological types have been described in prostate cancer, but we lack a clear understanding of the molecular mechanisms underlying these entities. It is likely that such causal mechanisms act at the DNA, RNA or protein levels. In this study, we identified several genomic changes (SNVs and CNAs) associated with a change in histological appearance. In all four instances of within-patient histological differences that we identified in this study, hundreds of unique SNVs place the distinct morphologies on separate branches of the evolutionary trees. Hence, histological heterogeneity in prostate cancer could represent later stages of divergent evolution. Our results also corroborate the previously reported finding that divergent lineages spatially coexist within the prostate^11^. Early work by Haffner et al. demonstrated the wealth of insights that can be drawn from multi-sample analysis in a single patient, identifying a histologically low grade prostate region as the source of the lethal metastatic clone^23^. Morphological features have been linked to molecular subtypes in other cancers^24,25^ and paradigms exist to combine both to predict patient outcomes^26^. In prostate cancer, specific morphological subtypes such as the cribriform pattern and intraductal carcinoma of the prostate have been linked with adverse outcomes^27,28^. Hence, our findings provide a strong case for histopathologically guided genomic studies to identify specific mutations associated with distinct morphologies at the cellular or glandular level, as well as to develop diagnostic methods that combine histological and genomic data.

Conversely, not all mutations are associated with a change in morphology; a sample from patient #10 annotated as ‘benign’ histologically was found to harbour a cluster of ∼250 SNVs representing a mutant clone. Cooper et al.^8^ and Erickson et al.^29^, demonstrated previously the presence of SNVs and copy number changes in histologically normal regions. Such regions may represent foci of early clonal expansion from a lineage distinct from the main tumour.

Taken together, we successfully integrated whole genome sequencing with histopathological data to unravel the source of intra-tumour histological and clonal heterogeneity in locally advanced prostate cancer. Using this approach, we obtained several new biological insights including identification of the genomic mechanism for the origin of amphicrine carcinoma of the prostate, and construction of detailed maps of intra-prostatic tumour evolution. We showed that morphologically distinct regions of cancer represent separate branches of evolution, which has implications for diagnosis. These insights will pave the way to develop novel tools to identify features of the prostate cancer metastatic and lethal phenotype, which in turn will allow early interventions where necessary and improved patient outcomes.

## METHODS

### Sample acquisition

Fresh frozen prostate samples were acquired from surgically resected (radical prostatectomy) specimens using a previously published method^30^. Lymph node and seminal vesicle metastases were additionally sampled when these structures were macroscopically involved by tumour, and care was taken that diagnostic processes would not be affected by biobanking. All samples were snap frozen in liquid nitrogen and stored at -80°C. After biobanking, specimens were processed for routine diagnostics by fixation in neutral buffered formalin for approximately 24 hours. Whole blood was collected at the time of surgery for germline DNA and stored at -80°C. Formalin-fixed, paraffin embedded samples were obtained through the routine histopathological diagnostic pathway.

### Sample processing

Serial frozen sections were cut from OCT embedded tissue and every 4^th^ section was stained for H&E and analysed tumour content by a specialist urological pathologist. Frozen sections were stored again at -80°C until nucleic acids extraction. DNA and RNA were extracted from the frozen sections using the Quick-DNA/RNA Miniprep Plus Kit (Zymo, D7003) according to the manufacturer’s recommendations. Briefly, 1X DNA/RNA Shield + proteinase K was added to the frozen sections and incubated at 50°C for 3 hours, followed by lysis and column extraction. Germline DNA was extracted from whole blood, also using the Quick-DNA/RNA Miniprep Plus Kit as above. DNA was quantified using a Qubit HS dsDNA kit (Thermo Scientific) and quality assessed using a TapeStation (Agilent Technologies).

### Whole Genome Sequencing and data pre-processing

DNA libraries were prepared and sequenced as 150bp paired-end reads by the Oxford Genomics Centre, Wellcome Centre for Human Genetics, Oxford on the Illumina NovaSeq 6000 using a PCR-free library preparation protocol or all samples except one (details in supplementary data - sample_list.csv). PCR amplification was performed for the sample with DNA quantity insufficient for the PCR-free protocol. The median depth of sequencing was 88X for the tumour samples, and 33X for the germline samples respectively. DNA libraries were prepared using the NEBNext Ultra II DNA Library Prep Kit for Illumina (New England Biolabs #E7645L) according to the manufacturer’s recommendations. Briefly, genomic DNA was fragmented and the ends were repaired enzymatically. Following this, adaptors were ligated, and the resulting library was cleaned up with NEBNext Sample Purification Beads. These libraries were used for sequencing in the PCR-free protocol. For samples requiring amplification, these libraries were PCR-amplified and cleaned up using NEBNext Sample Purification Beads.

### Sample processing and library preparation for FFPE samples

Pathologist-marked regions of interest were manually dissected and pooled from 5X serial sections (4 um) of formalin fixed paraffin embedded radical prostatectomy samples. These dissected samples were then used for an extraction-free WGS library preparation method modified from Ellis et al.^31^. Briefly, samples were manually dissected using histology-guided markings and pooled from 5 × 4 μm FFPE sections. The samples were deparaffinised and lysed in a lysis buffer (Tris-HCl: pH 8.0 30mM, Tween-20: 0.5%, IGEPAL CA-630: 0.5%) containing Proteinase K (20mg/ml) and incubated at 65°C on a heating block for 2-3 hours. Following lysis, the samples were size selected using Ampure XP Beads and fragmented and end-repaired for 12 minutes using the NEB Ultra II FS enzyme. The end-repaired DNA samples were ligated with custom adapters described in Ellis et al. and amplified with single-indexed primers to generate WGS libraries for Illumina sequencing. The individual libraries were pooled and sequenced as 150bp paired-end reads on the Illumina NovaSeq 6000 sequencer at Wales Gene Park, Cardiff. The median depth of sequencing for the FFPE samples was 60X.

### Data pre-processing

Raw data in the form of FASTQ files were adapter-trimmed using BBDuk^32^ (v38.79-0) and pre- and post-trim quality was checked using FastQC^33^ (v0.11.8). Reads were aligned to the reference genome (hg37) using BWA-MEM^34^ (v0.7.17) and coordinate-sorted using Samtools^35^ (v1.9). Aligned reads from all samples (PCR-free as well as PCR-amplified) were duplicate marked using Genome Analysis ToolKit (GATK)^36^ (v4.1.4.0) MarkDuplicates.

### SNV calling

Base Quality Score Recalibration (BQSR) was performed on duplicate marked bam files and simple somatic mutations (SNVs and short indels) calling was performed with GATK Mutect2 in multitumour mode (i.e., by calling somatic mutations in all samples from a patient simultaneously) according to the GATK Best Practices Workflow. Calls from Mutect2 were filtered using GATK FilterMutectCalls. SNVs were further shortlisted for subclone analysis by using a TLOD (log likelihood of a variant being somatic) threshold of 7.

### CNA calling

Allele-specific subclonal copy number calling was performed using the Battenberg R package^37^ (v2.2.9) from the base quality recalibrated, duplicated marked bam files. Default parameters were used initially to generate copy number profiles. These were used to run subclonal analysis as follows, and the resultant CCF values for each SNV were plotted along the genome (Supplemental information) as a quality check. Where regions of the genome were found to contain errors in CCFs (CCFs less than surrounding regions in case of incorrect calls of amplification/gain suggesting overcorrection, or CCFs greater than the surrounding regions in case of incorrect calls of LOH/deletion suggesting undercorrection). In addition, the expected purity of the sample was estimated by adjusting the CCF of the truncal cluster to ∼1. Where the original CNA calls were found to be incorrect using these two checks, Battenberg was run again with rho and ploidy presets and checked again for accuracy iteratively. In cases where the tumour purity was too low to call CNAs with Battenberg (lymph nodes 5A_LN, 6A_LN, 7A_LN in patient #02, PPF_LN in patient #13), the most common CNA profile from the intra-prostatic samples was applied, and a quality check was performed as above to confirm that this is a reasonable assumption to make. Percent subclonality was calculated by dividing the total number of bases with subclonal copy number changes with the total number of bases analysed.

### Phylogenetic analysis

The DPClust-3p R package^38^ (v1.0.8) was used to pre-process the SNVs (Mutect2) and copy number calls (Battenberg) from each sample and generate loci with CCF (allele fraction corrected by the corresponding copy number for that location). Following this, the CCFs of the loci were clustered using the BayesianGaussianMixture function in the Scikit-Learn Python package^39^ (v0.24.2) using 1000 iterations. The median value of each cluster was considered to be the corresponding clonal fraction for that subclone. Clusters thus identified were further split or merged by visual inspection where deemed necessary (e.g. when a cluster was visually determined to be composed of multiple sub-clusters). From the CCFs of clones thus obtained, phylogenetic trees were constructed using the sum and crossing rules as described previously^40^. Briefly, the sum rule states that the sum of CCFs of daughter clones should be less than the CCF of the parent clone. If this is not the case, the daughter clone with the smaller number of mutations is a descendant of the other daughter clone. The crossing rule, applicable in multi-region sequencing, states that when clones B and C are daughter clones of clone A, and CCF(B) > CCF(C) in one sample and CCF(C) > CCF(B) in another sample, B and C must be clones on different branches of the phylogenetic tree. Where the sum and crossing rules contradicted each other, a solution that violated the fewest rules was chosen, to allow for inherently noisy biological data.

### Construction of clone maps

Clonal composition of each tumour sample was illustrated as a sunburst plot using Plotly^41^ (v5.3.1). Corrections were applied to the CCFs based on the solution calculated, as described above, to simplify visualisation. Actual CCFs are presented in the supplementary data (ccf_tables.xlsx).

### Mutation Signature analysis

Mutational signatures were identified using the DeconstructSigs^42^ (v1.8.0) R package. SNVs were grouped into separate bins based on their cluster annotation and each group was processed as a separate ‘sample’. COSMIC v3 trinucleotide signatures were used as the reference.

### Structural Variant analysis

Structural variants were called using Svaba^43^ (v1.1.0) using default parameters in multi-tumour mode.

### Software

All analyses were performed in R^44^ (v4.2.1) and Python^45^ (v.3.9.6).

### Database sources

Prostate cancer specific driver mutations were downloaded from Intogen^46^.

### Immunohistochemistry staining

#### PTEN, P-Histone H3 (PHH3) Immunohistochemistry (IHC)

PTEN and PHH3 IHC was performed on a Bond RX autostainer (Leica Biosystems) using a 2 day protocol. On day 1, slides of formalin-fixed paraffin-embedded sections were loaded into the Bond Rx autostainer. The slides were deparaffinised with Leica Dewax solution (AR9222). Epitope Retrieval was performed using Solution 1 (AR9961): 30 minutes for PTEN IHC and 20 minutes for P-Histone H3 IHC. Endogenous peroxidase was blocked using (Bond Polymer Refine Detection Kit DS9800). The slides were unloaded from the Bond Rx. For PTEN IHC, 100 micro litres of PTEN Rabbit monoclonal antibody (Cell Signaling 138G6) diluted 1/200 in Leica Antibody Diluent (AR9352) was applied to the sections. For PHH3 IHC, (Cell Signaling 9701S) diluted 1/100 in Leica Antibody Diluent (AR9352) was applied. The sections were incubated overnight at 4°C in a humidity chamber. On day 2, the slides were loaded into the Bond Rx autostainer and the post primary steps were completed using the Bond Polymer Refine Detection Kit (Leica DS9800). For Ki-67 staining, Leica RTU Ki67 PA0410 antibody was used at a dilution of 1:200, epitope retrieval for 20 minutes, using the JR Bond protocol F + enhancer (includes 15 minutes antibody incubation). Scoring of PTEN IHC was done by a board certified pathologist (CV) using H-Scoring^47^ which involves assessment of intensity (0-3) and proportion score (0-100%).

#### PSMA Immunohistochemistry

Formalin-fixed paraffin-embedded sections were heated to 60°C for 10 minutes, deparaffinised in xylene and rehydrated in graded concentrations of ethanol. Endogenous peroxidase was neutralised with 3% H_2_O_2_ in methanol for 10 min at room temperature. Antigen was retrieved using citrate buffer pH 6 for 10min at 95°C and samples were left to cool in the buffer for 20 minutes. Samples were blocked with PBS/5% NGS for 30 min at room temperature. Mouse PSMA monoclonal antibody (Gene Tex GTX19071) diluted 1/100 in PBS/5% NGS was applied to the samples overnight at 4 o C. The sections were then incubated with Biotinylated goat anti mouse IgG (Vectorlabs BA-9200) diluted 1/250 in PBS/5% NGS for 30 min at room temperature. The detection system was ABC reagent (Vectorlabs PK-7100) and DAB Substrate Kit SK-4100 (Vectorlabs). Counterstain was Harris’s haematoxylin.

### Image analysis

Stained slides were scanned on the Hamamatsu scanner and analysed using the VisioPharm image analysis platform. Different regions of tumour were annotated by the pathologist (CV). A small subset of immune (H&E) or positive staining cells (IHC) cells were manually selected for training a model which was then used to count the number of immune cells in each annotated region.

## Data and code availability

Sequencing data is being deposited in the European Genome-Phenome archive and all code used to generate these figures will be deposited in a GitHub repository.

## Acknowledgements

We are grateful to Cancer Research UK (C1380/A18444) and NIHR Oxford BRC Surgical Innovation Theme for funding this research. We acknowledge the contribution to this study made by the Oxford Centre for Histopathology Research and the Oxford Radcliffe Biobank. We thank the Wellcome Trust Centre for Human Genetics, Oxford and Wales Gene Park, Cardiff for sequencing services. The computational aspects of this research were funded from the NIHR Oxford BRC with additional support from the Wellcome Trust Core Award Grant Number 203141/Z/16/Z. The views expressed are those of the author(s) and not necessarily those of the NHS, the NIHR or the Department of Health.

## Author contributions

SR, LC, ZK, MJ performed laboratory experiments. SR, CV, DJW, NKA, FB, DCW generated and analysed data. SR, CV, DJW interpreted the results. CV, MO’H, AH, AL, curated and biobanked patient samples. ADL, ITul, JN, SL, AO, FL, TL were involved in surgical sample collection. FH, IT and BV procured funding and supervised the project. SR wrote the manuscript with input from CV, DJW, FH, IT, IM, AL, RB.

