## Supplemental figures for "Intra-prostatic tumour evolution, steps in metastatic spread and histogenomic associations revealed by integration of multi-region whole genome sequencing with histopathological features"

1. **Pairwise plots of SNV CCFs**

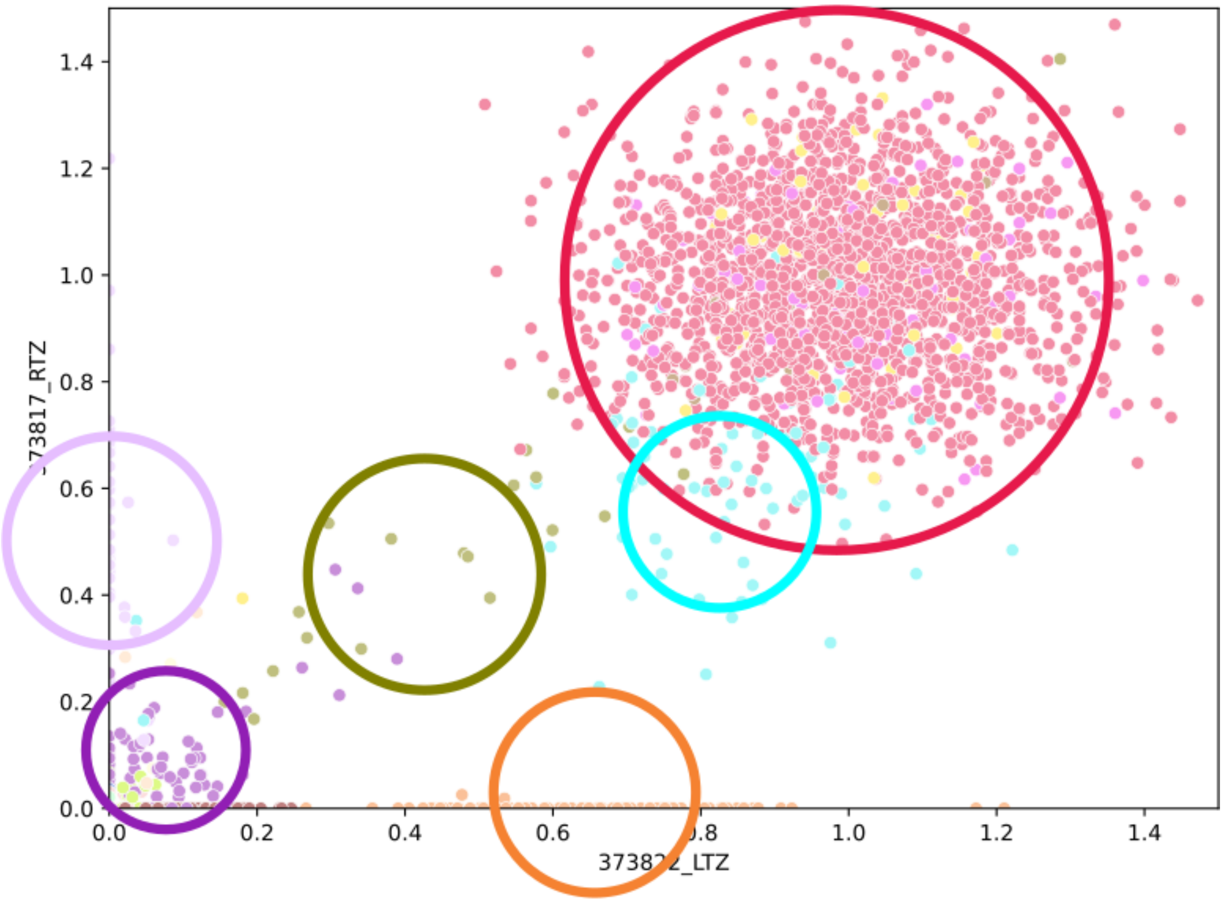


Figure S1. Pairwise plots of SNV CCFs were used to visually confirm accurate calling of clusters. The median CCF value of each cluster was taken as the CCF of the subclone. The CCF of the truncal cluster must be 1.

2. **Patient #02**

Copy number profile

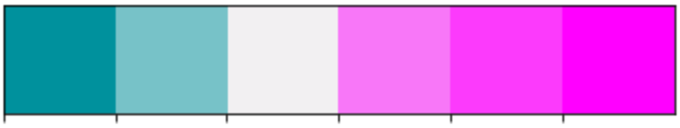

Figure S2. Copy number profile plots are colour-coded as above, with dark and light blue representing homozygous (0) and heterozygous (1) deletions respectively, grey denoting a diploid copy number (2), and shades of pink representing increasing total copy number counts (>= 3).

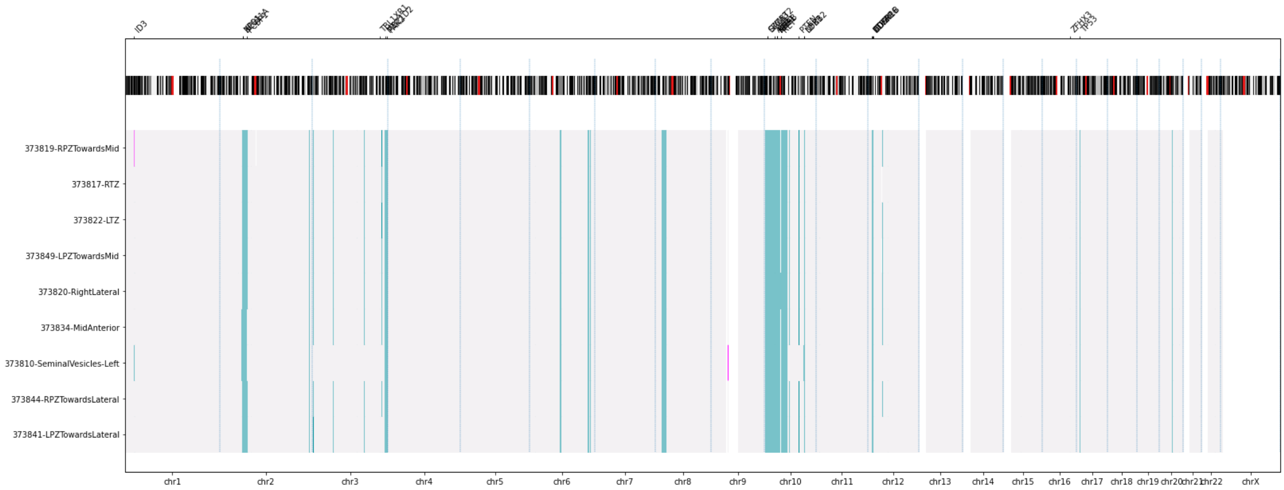


Figure S3. Copy number profile for patient #02, with genomic coordinates along the X axis and sample names along the Y axis (bottom). Cytoband data for each chromosome (to help with visualisation of centromeres which are depicted as red stripes) and known driver genes present in the CNAs are annotated at the top of the plot. Lymph node samples 5A, 6A and 7A were assumed to have the same profiles as the majority of the intra-prostatic samples; hence the copy number profiles for these samples are not shown.

### CCF-cluster plot

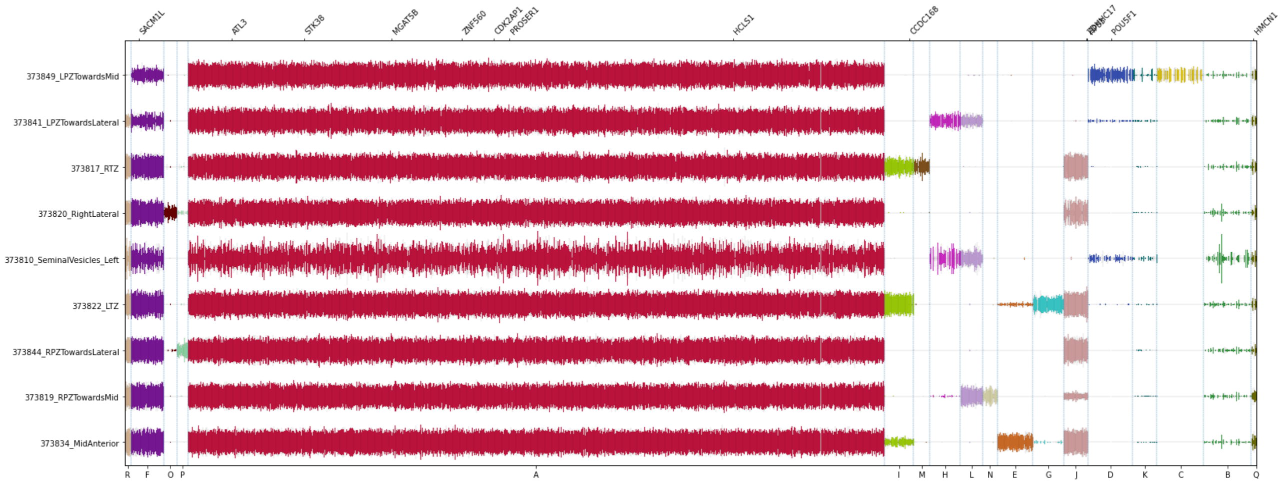


Figure S4. In the CCF-cluster plot, SNVs represented as vertical lines are grouped by their cluster annotation along the X axis (bottom) and sample names are along the Y axis. The vertical length of each SNV is proportional to its copy number adjusted CCF. Missense mutations in known coding regions are annotated along the top of the X axis. Low tumour purity samples (e.g. 373810_SeminalVesicles_Left) are more noisy. SNVs from clusters K, B, and Q cannot be placed on the phylogenetic tree and may represent noise. Hence these SNVs were ignored.

### 3. **Patient #08**

Copy number profile

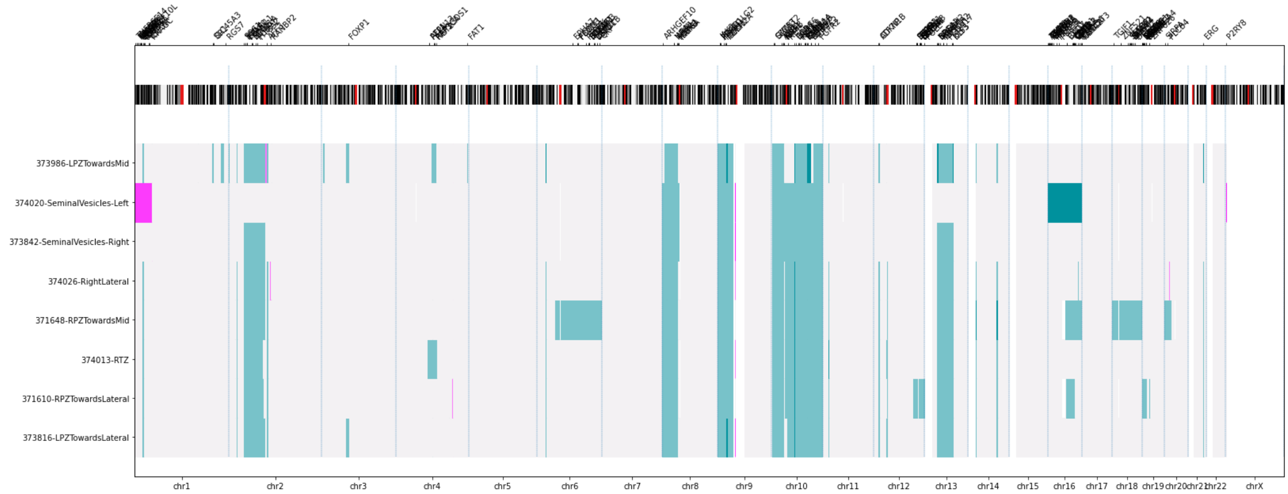


Figure S5. 374020-SeminalVesicles_Left is of lower tumour purity (0.06) and some CNA calls (e.g. 1p amplification, 16 deletion, and missed CNA calls 2p LOH, 13q LOH) are inaccurate.

CCF-cluster plot

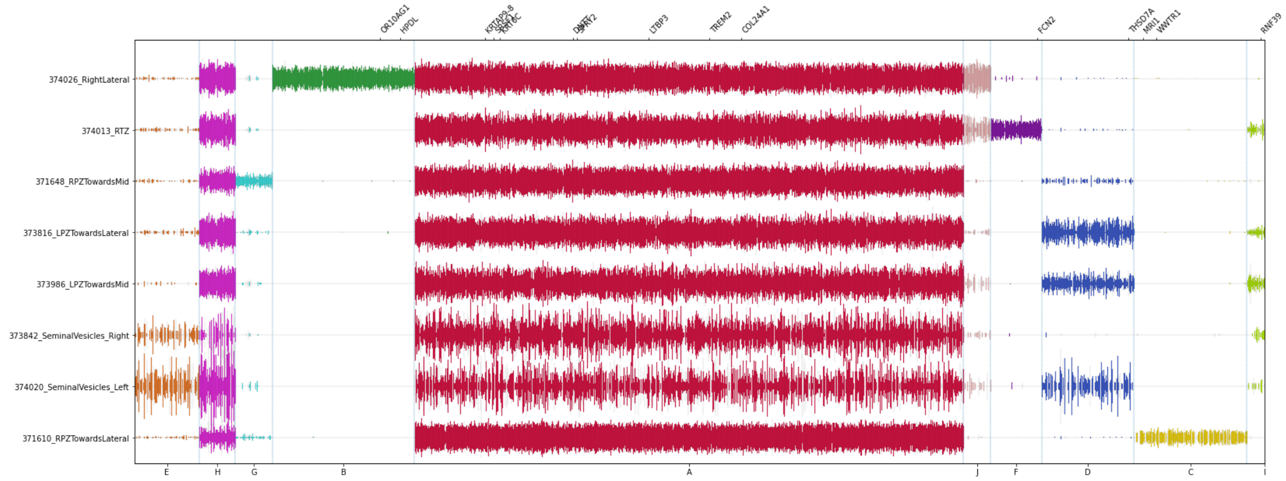


Figure S6. SNVs from cluster I cannot be placed on the phylogenetic tree and may represent noise. Hence these SNVs were ignored.

4. **Patient #13**

Copy number profile

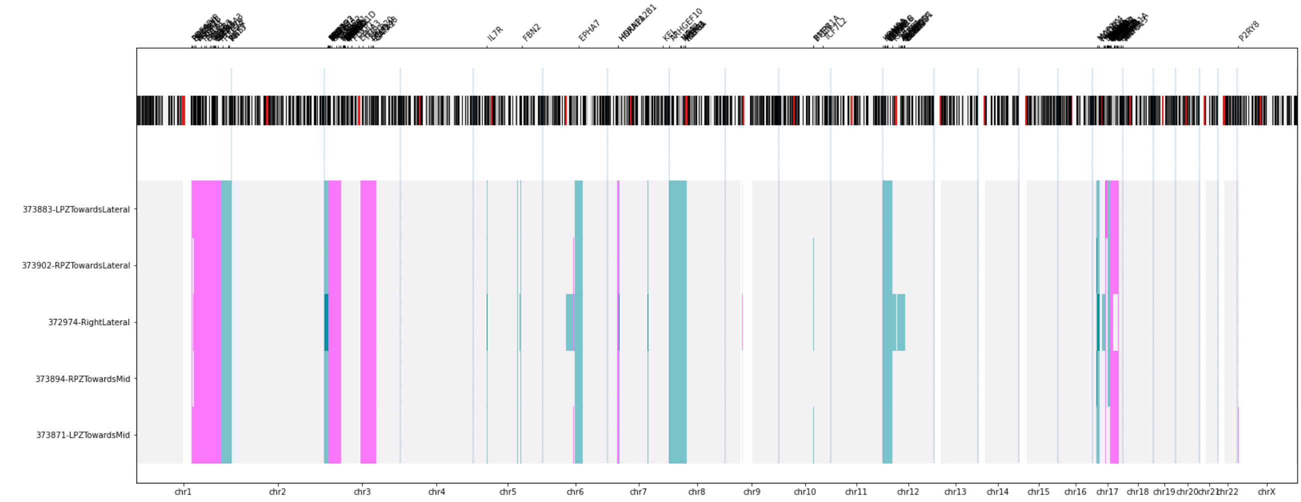


Figure S7. 373901-Pre-prostaticFat is of lower tumour purity (0.08) and hence was assumed to have the same CNA profiles as the majority of the intra-prostatic samples; the copy number profiles for these samples are not shown.

CCF-cluster plot

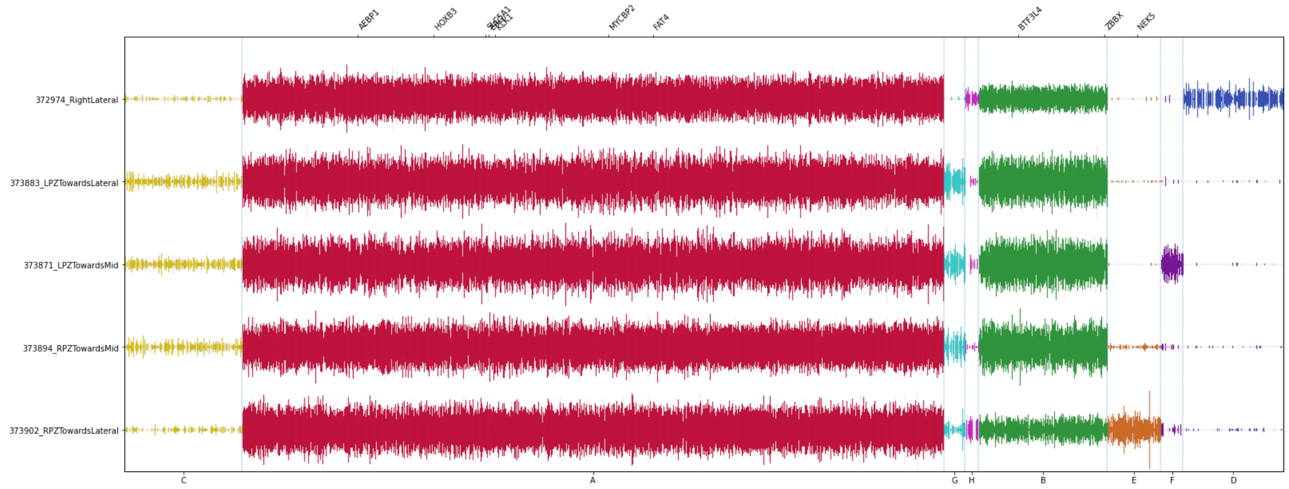


Figure S8. SNVs from clusters G and H cannot be placed on the phylogenetic tree and may represent noise. Hence these SNVs were ignored.

5. **Patient #10**

Copy number profile

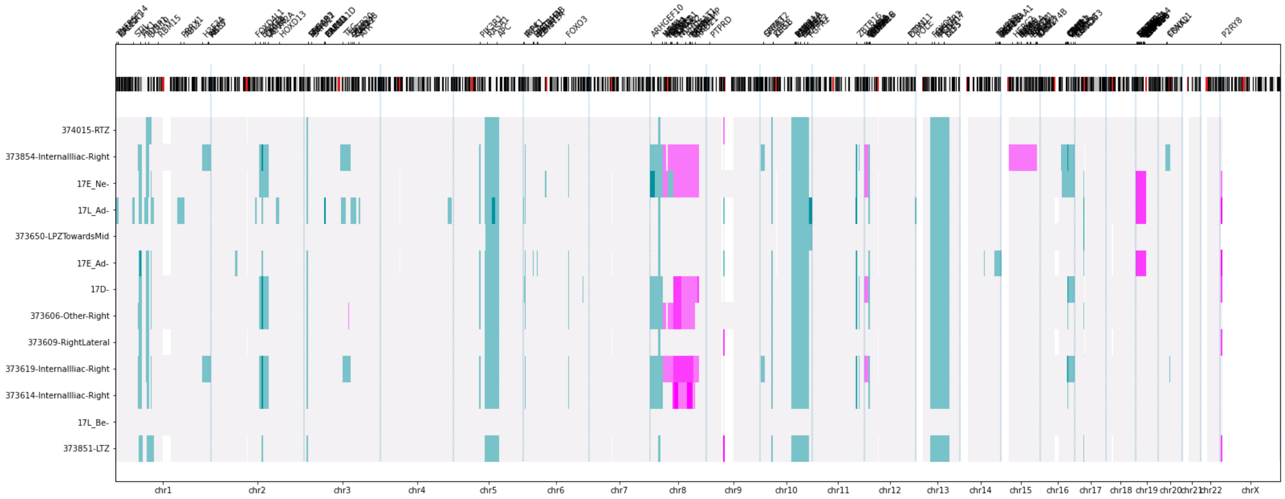


Figure S9. Amplifications in 8p are seen in the lymph node samples and intra-prostatic neuroendocrine samples.

CCF-cluster plot

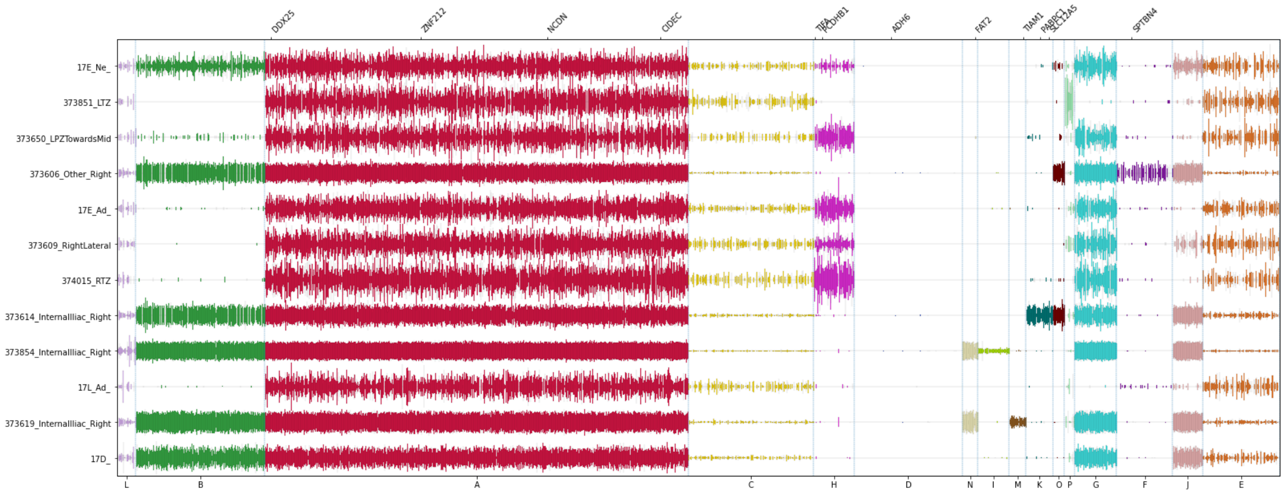


Figure S10. SNVs from clusters L, C, P, E cannot be placed on the phylogenetic tree and may represent noise. Hence these SNVs were ignored. Cluster D is present in the MidAnterior sample but not in any other samples, suggesting a distinct lineage.

6. **Patient #15**

Copy number profile

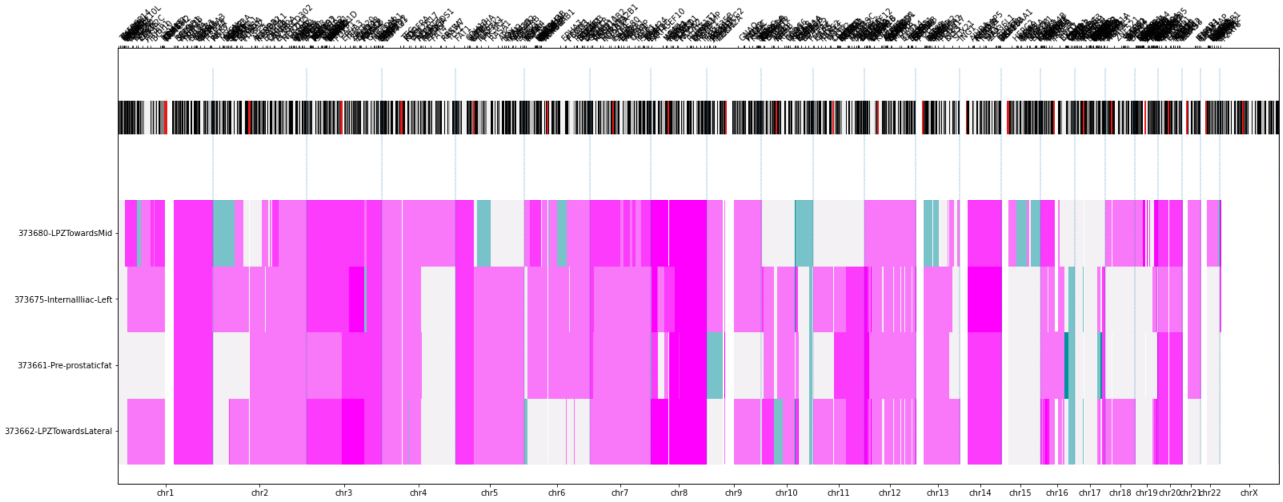


Figure S11. Extensive copy number alteration is seen in all samples from this patient.

CCF-cluster plot

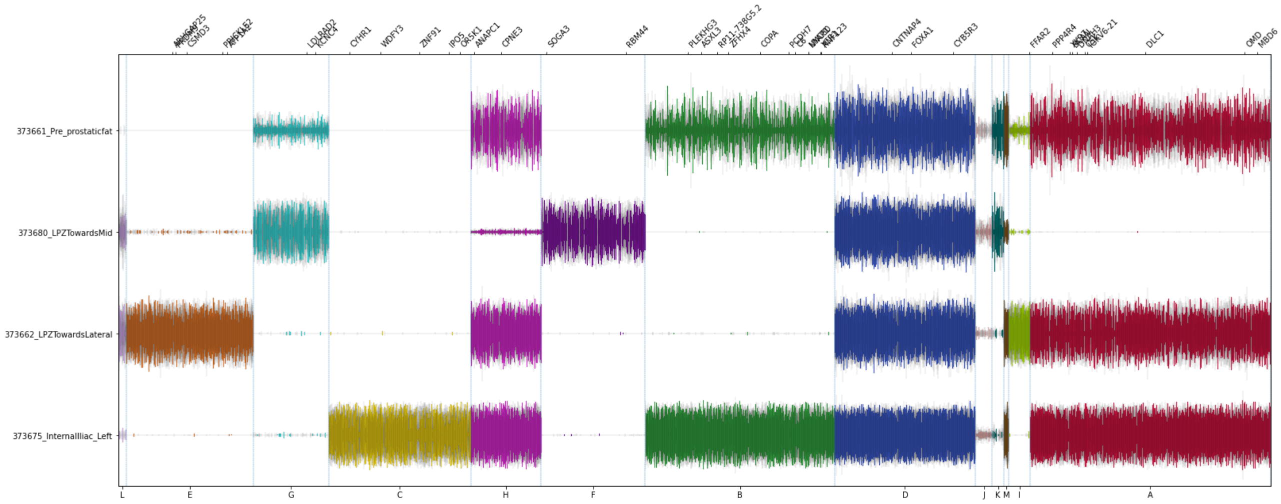


Figure S12. SNVs from clusters L, J, K, M cannot be placed on the phylogenetic tree and may represent noise. Hence these SNVs were ignored.

7. **Knockdown of FOXP1 results in increased migration in vitro**

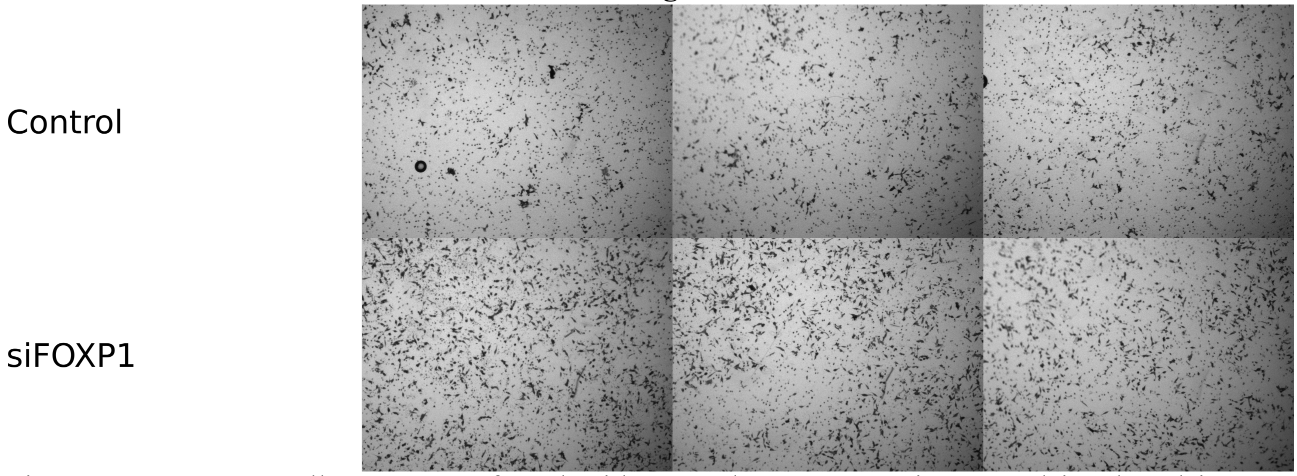


Figure S13. LNCaP cells were transfected with control or FOXP1 siRNA and incubated in 8um transwells for 48 hours. Cells with FOXP1 knockdown have increased migration compared to control cells.

8. **Patient #15: BRCA CN status**

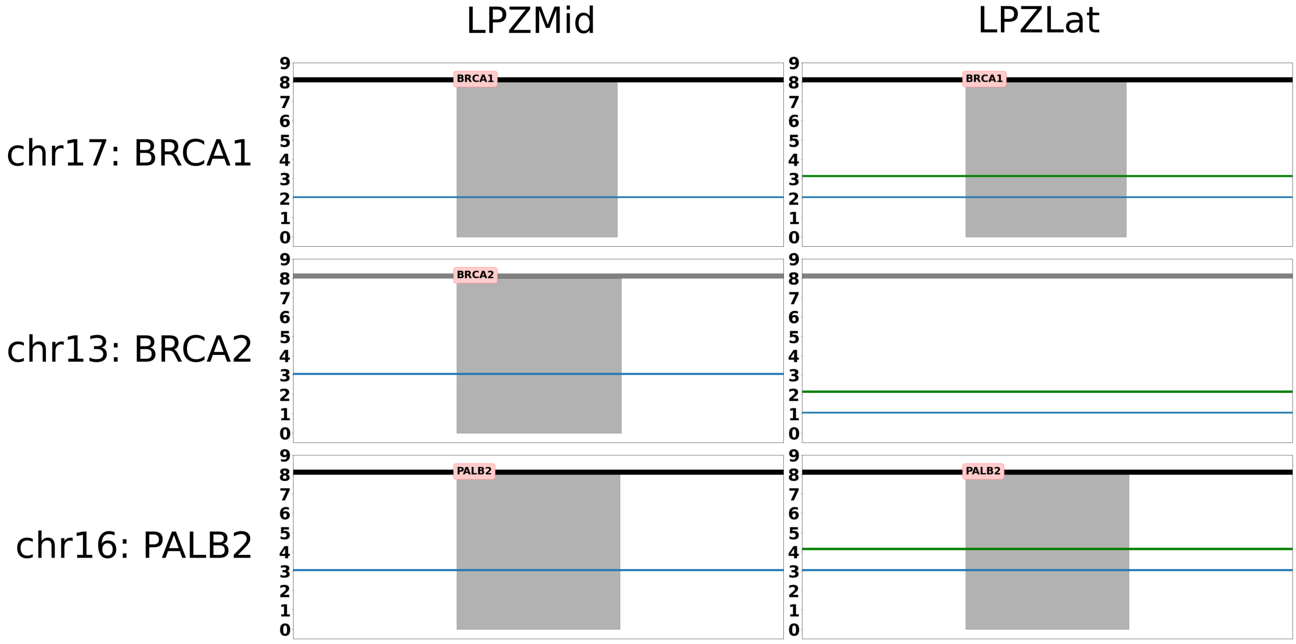


Figure S14. Copy number status at BRCA1, BRCA2 and PALB2 loci, derived from Battenberg calls, are plotted for LPZMid and LPZLat samples. Blue line indicates major copy number and green line indicates minor copy number.

9. **Patient #15: BRCA methylation status**

BRCA1

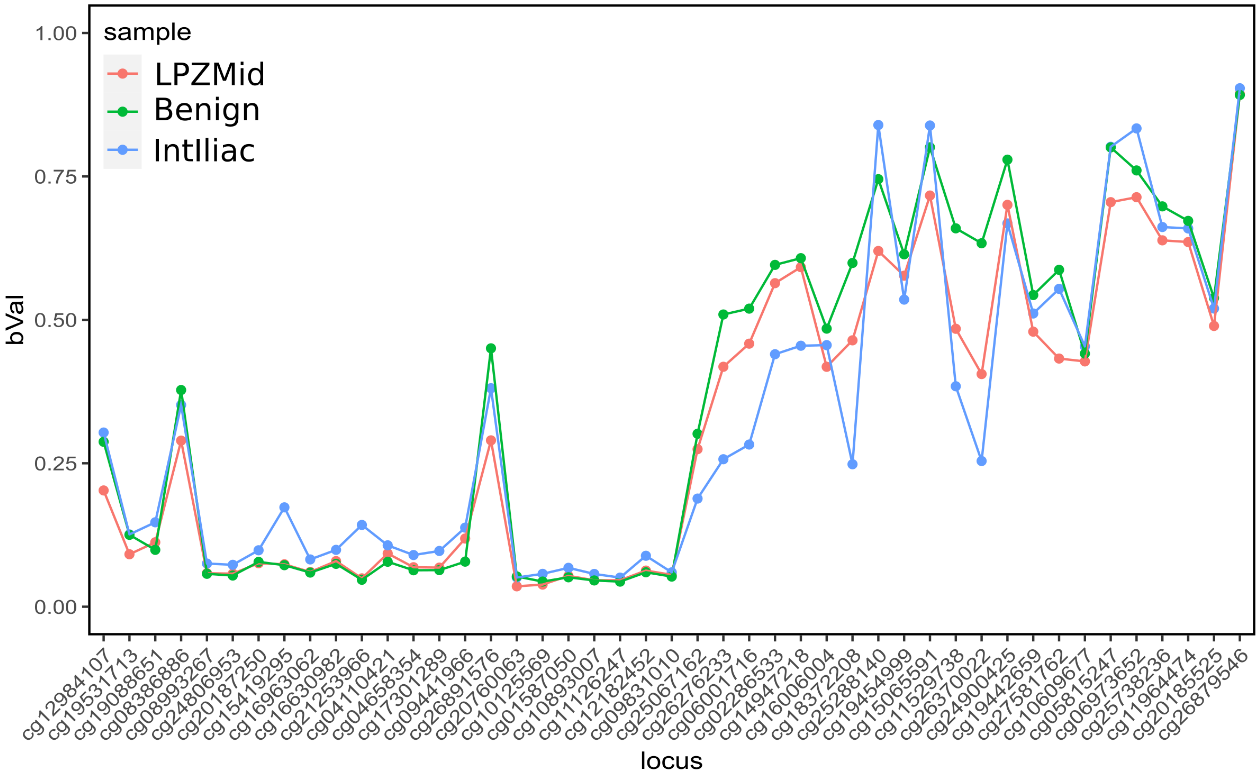


Figure S15. Methylation levels in the BRCA1 promoter region are plotted in 3 samples from patient #15.

BRCA2

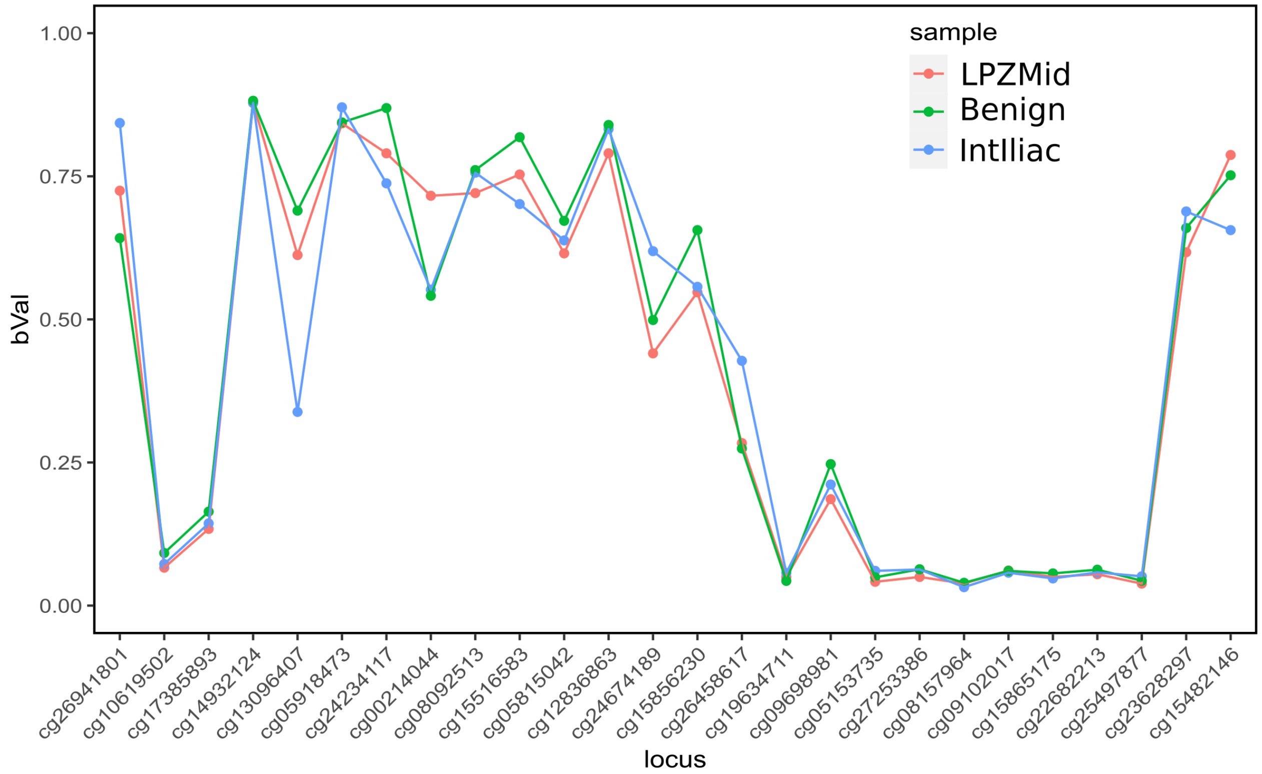


Figure S16. Methylation levels in the BRCA1 promoter region are plotted in 3 samples from patient #15.

10. **Adenocarcinoma in #10**

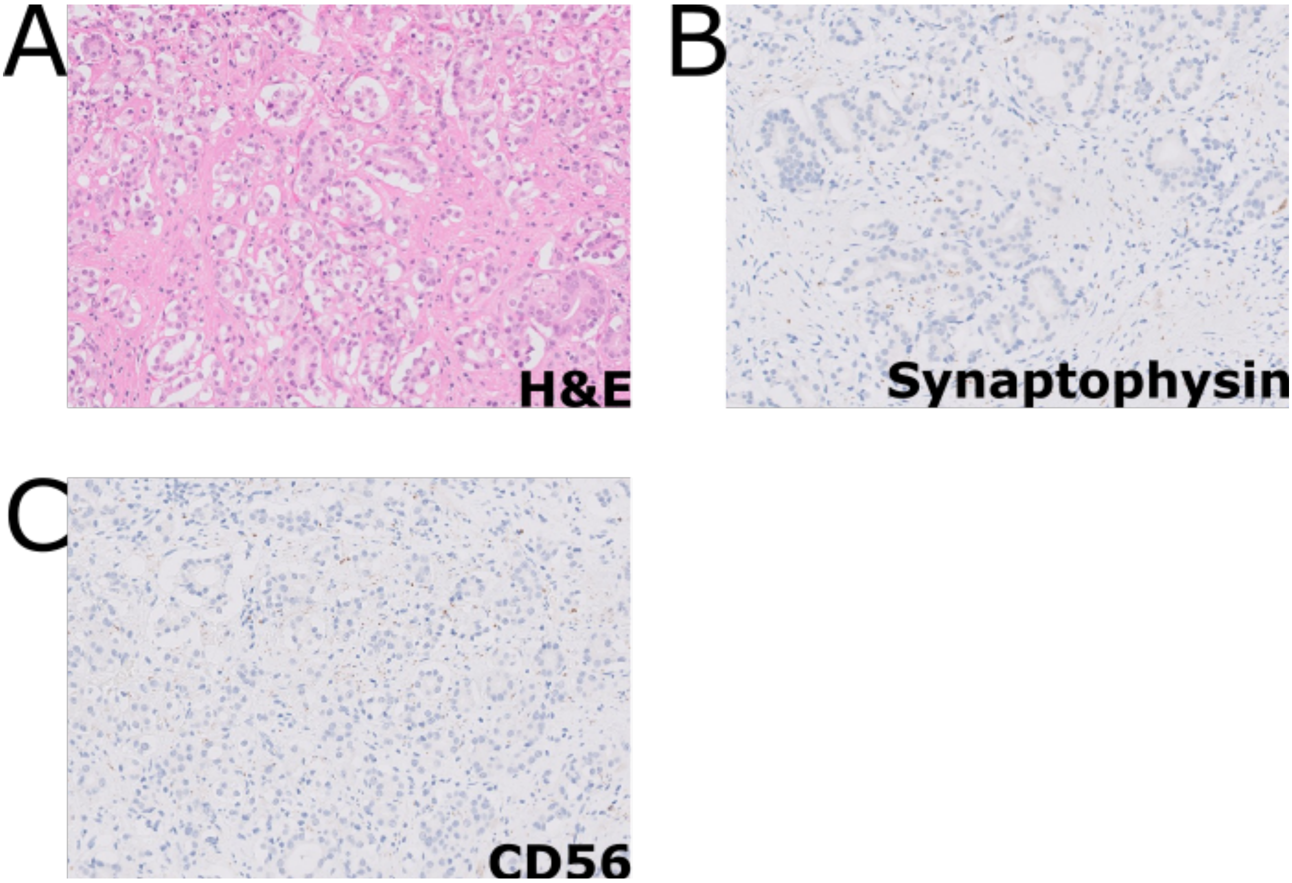


Figure S17. The intra-prostatic tumour in patient #10 with an acinar adenocarcinoma morphology as seen by H&E (A) stained negative for the neuroendocrine markers Synaptophysin (B) and CD56 (C).
